# Identification of a novel LysR-type transcriptional regulator in *Staphylococcus aureus* that is crucial for secondary tissue colonization during metastatic bloodstream infection

**DOI:** 10.1101/2020.06.23.166322

**Authors:** Michaela Groma, Sarah Horst, Sudip Das, Bruno Huettel, Maximilian Klepsch, Thomas Rudel, Eva Medina, Martin Fraunholz

## Abstract

*Staphylococcus aureus* is a common cause of bacteremia that can lead to severe complications once the bacteria exit the bloodstream and establish infection into secondary organs. Despite its clinical relevance, little is known about the bacterial factors facilitating the development of these metastatic infections. Here, we used a *S. aureus* transposon mutant library coupled to transposon insertion sequencing (Tn-Seq) to identify genes that are critical for efficient bacterial colonization of secondary organs in a murine model of metastatic bloodstream infection. Our transposon screen identified a LysR-type transcriptional regulator (LTTR), which was required for efficient colonization of secondary organs such as the kidneys in infected mice. The critical role of LTTR in secondary organ colonization was confirmed using an isogenic mutant deficient in the expression of LTTR. To identify the set of genes controlled by LTTR, we used a *S. aureus* strain carrying the LTTR gene in an inducible expression plasmid. Gene expression analysis upon induction of LTTR showed increased transcription of genes involved in branched chain amino-acid biosynthesis, a methionine sulfoxide reductase and a copper transporter as wells as decreased transcription of genes coding for urease and components of pyrimidine nucleotides. Furthermore, we show that transcription of LTTR is repressed by glucose, induced under microaerobic conditions, and required trace amounts of copper ions. Our data thus pinpoints LTTR as an important element that enable a rapid adaptation of *S. aureus* to the changing host microenvironment.

## Introduction

*Staphylococcus aureus* is a common colonizer of the human skin and mucosal membranes without causing diseases. However, *S. aureus* can also cause a variety of severe infections such as sepsis, abscesses in deep tissues and osteomyelitis after reaching sterile anatomic sites for example via the blood stream (1). The emergence and spread of *S. aureus* strains resistant to multiple antibiotics (2) added to the remarkable capacity of *S. aureus* to evade elimination by the host immune defenses (3) makes this pathogen a formidable challenge for physicians. Despite extensive efforts made by the research community to uncover the pathogenic strategies of *S. aureus*, many aspects of the infection process remain unclear. In metastatic bloodstream infections, *S. aureus* disseminates from a primary site of infection such as the skin abscesses, intravenous catheter or surgical sites to secondary organs (4). Several virulence factors have been reported to mediate the interaction and/or extravasation of *S. aureus* through the endothelium including the extracellular adherent protein Eap, wall teichoic acid or fibronectin binding proteins (5). After extravasation from the bloodstream to adjacent tissue, *S. aureus* must adapt to the physicochemical microenvironment, nutrient availability as well as to the local immune response in the new niches in order to survive, proliferate and establish productive infection. The virulence mechanisms of *S. aureus* that facilitate these processes remain largely unknown. A deeper understanding of these pathogenic mechanisms can provide novel opportunities for therapeutic interventions in bacteremic patients.

We have previously reported the remarkable plasticity of *S. aureus* to reprogram its expression of virulence factors during infection to adapt to the microenvironment and biological pressure imposed by the host (6, 7). In this study, we have used a *S. aureus* transposon insertion library coupled to deep sequencing of transposon insertion sites (Tn-Seq) to identify genes required by *S. aureus* to colonize and establish infection in secondary organs during in vivo infection. In general, Tn-Seq has become a very popular high throughput technique that has been widely applied to identify genes and pathways that are important for infection for different pathogens (8–10). We and others have successfully applied this technique to identify novel factors involved in *S. aureus* pathogenicity (11, 12). Thus, in a previous study using a transposon mutant library generated with the staphylococcal strain 6850 we identified the araC-type transcriptional regulator repressor of surface proteins rsp as a pleiotropic virulence factors regulator during the initial stages of infection (11). Here, we screened this *S. aureus* transposon insertion library (11), in a well-established mouse model of metastatic bloodstream infection (13). Mice were infected with the mutant library and the mutants recovered from the primary site of infection, the liver, as well as from the secondary infected organs, the kidneys in this case, were compared with those present in the original inoculum. The mutants underrepresented in the pool of colonizing bacteria in comparison with the original inoculum indicated the genetic determinants that are critical for optimal establishment of infection in secondary organs. A single gene was identified (RSAU_000852) that encoded for a LysR-type transcriptional regulator (LTTR), whose mutants were specifically depleted in the bacteria pool recovered from the kidneys. An isogenic mutant in LTTR exhibited significantly reduced capacity to survive in kidneys and tibiae than the wild type, corroborating thus the results of the Tn-Seq analysis. Functional characterization of LTTR indicated that this regulator may be required by *S. aureus* to rapidly adjust to the environmental conditions and nutrient availability in secondary organs.

## Materials and Methods

### Mice and infection model

Pathogen-free 10 weeks old female C57BL/6 mice (20.5±0.8 g body weight) were purchased from Harlan-Winkelmann (Envigo, The Netherlands). Mice were infected intravenously with 10^6^ colony forming units (CFU) of *S. aureus* strain 6850 in 100ml of PBS via a lateral tail vein. For determination of bacterial numbers in organs, mice were killed by CO_2_ asphyxiation at the specified time after bacterial inoculation, organs were removed and homogenized in PBS. Serial 10-fold dilutions of organ homogenates were plated on blood agar plates. Bacterial colonies were counted after incubation at 37°C for 24 h and calculated as CFU per organ. Animal experiments were performed in strict accordance with the German regulations of the Society for Laboratory Animal Science (GV-SOLAS) and the European Health Law of the Federation of Laboratory Animal Science Associations (FELASA) and animals were excluded from further analysis if killing was necessary according to the human endpoints established by the ethical board. All experiments were approved by the ethical board Niedersächsisches Landesamt für Verbraucherschutz und Lebensmittelsicherheit, Oldenburg, Germany (LAVES; permit N. 33.9-42502-04-13/1195).

### ELISA

The concentration of IL-6 in serum of infected mice was determined by ELISA according to the manufacturer’s recommendations (BD Biosciences).

### Bacterial culture

*S. aureus* strains (Table S1) were grown in trypticase soy broth (TSB), lysogeny broth (LB) or chemically defined medium (CDM) using appropriate antibiotics. The TnSeq inoculum was prepared by growing the strains in brain-heart infusion (BHI) to a OD of 0.5, washed with PBS, aliquoted and stored at −80°C.

CDM contained final concentrations of 1x basic medium (12.5 mM Na_2_HPO_4_ 2H_2_O, 10 mM KH_2_PO_4_, 1.65 mM MgSO_4_, 9.25 mM NH_4_Cl, 8.5 mM NaCl); 75 mM glucose; 0.142 mM sodium citrate tribasic dihydrate; 1 mM of each amino acid, except for tyrosine (0.1 mM). The trace element mix A5 was purchased from Sigma-Aldrich (#92949) and was supplemented with FeCl_3_ in NaOH; 0.1 μM NiCl_2._ Vitamin mix contained final concentrations of 0.29 μM 4-Aminobenzoic acid, 0.29 μM Thiamine hydrochloride, 0.21 μM Ca-D-Pathothenic acid, 0.036 μM Cyanocobalamin, 0.04 μM D-Biotin, 0.81 μM Nicotinamid, 0.62 μM Pyridoxine hydrochlorid, 0.2 μM Riboflavin.

For induction of LTTR expression by anhydrous tetracycline, overnight cultures were diluted in 10 ml fresh TSB containing 200 ng/ml anhydrous tetracycline and were incubated at 37° C, 750 rpm for 1 h. Bacteria were harvested and snap-frozen in liquid nitrogen.

### Promoter activity assays

Promoter activities during bacterial growth were assessed by monitoring GFP fluorescence (excitation, 488 ± 9 nm; emission, 518 ± 20 nm) as well as optical density (600 nm) using a Tecan Infinite M200 multi plate reader. For this, bacteria were grown in either TSB, LB, CDM or CDM without glucose overnight at 37°C, 180 rpm in the presence of 20 μg/ml chloramphenicol. If media exchange was necessary, overnight cultures where washed once with PBS and were resuspended in the corresponding media. Bacterial suspensions were added with an OD of 0.1 to a 48-well microtiter plate (400 μl/well) and further cultured 6 h or 24 h under aerobic and microaerobic conditions. A microaerobic atmosphere was established by sealing the microtiter plates with adhesive foil and thereby limiting oxygen availability. Microaerobic conditions were confirmed by the relative induction of the gene encoding formate acetyltransferase (*pfl*B), which is expressed only under low oxygen conditions (14). Absorbance and GFP fluorescence were recorded every 10 min using a Tecan Infinite M200 multi plate reader and were evaluated using Microsoft Excel.

### *Staphylococcus aureus* Tn-Seq

Pooled mariner transposon mutant libraries were generated as previously described (11, 15). Bacterial DNA was isolated from the infection inoculum and from the bacterial colonies derived from infected tissues as previously described (11). The libraries were sequenced on the Illumina^®^ Hi-Seq 2500 platform with the transposon-specific oligonucleotide primer Himar1-Seq (11). Illumina adapter sequences were removed via cutadapt version 1.2.1 (16). The reads also were filtered for size (>16 bp) and to contain the transposon ITR and were mapped to the *Staphylococcus aureus* 6850 genome (RefSeq accession NC_022222.1) by Bowtie2 v2.1.0 (17). Identification of depleted and enriched mutants was performed via DESeq2 version 1.6.2 (18).

### Cloning procedures

Chromosomal DNA of *S. aureus* was prepared using the QlAprep®Spin miniprep kit from Qiagen with the following modification: after resuspension of the bacteria in buffer P1 we added 50 μl lysostaphin and incubated at 37°C for 30 min (750 rpm).

For deletion of RSAU_000852 in *S. aureus* 6850, regions approximately 1 kb upstream and downstream were amplified by PCR using primers attB1-852-up-F and 852-up-R-SacII as well as 852-down-F-SacII and attB2-852-down-R, respectively (Table S2). The PCR product was cloned into pKOR1 and a marker-less targeted gene deletion was then generated as previously described (19). In order to exclude relevant secondary site mutations we sequenced the genome of the mutant (www.eurofinsgenomics.eu). Sequence reads were mapped to the genome sequence of the wild-type *S. aureus* 6850 genome (RefSeq accession NC_022222.1).

The RSAU_000852 ORF was amplified from *S. aureus* genomic DNA using oligonucleotides 852-NotI-F and 852-BamHI-R (Table S2). Plasmid p2085-852, allowing for anhydrous tetracycline-inducible expression of RSAU_000852 was generated by restricting the PCR product as well as the vector, p2085 (20), with BamHI and NotI and subsequent ligation of vector and insert. The resulting plasmid was transformed into E. coli DH5α and plated on LB-agar supplemented with ampicillin. The plasmid was isolated from E. coli was sequenced, propagated within the cloning strain *S. aureus* RN4220 and transformed into electro-competent *S. aureus* 6850.

For generation of the LTTR promoter (pr852) activity construct, p2085 was amplified using oligonucleotides pGFP-Inf-Prom-F and pGFP-vec-R (Table S2). pr852 was amplified from *S. aureus* DNA using oligonucleotides pGFP_852-Prom_fw and pGFP_852-Prom_rev. This fragment was cloned in the linearized vector using infusion cloning, was transformed into E. coli DH5α and plated on LB agar plates supplemented with 100 μg/ml ampicillin. The plasmid was isolated from E. coli, the insert sequence was verified by Sanger sequencing and electroporated into *S. aureus* RN4220 and finally transferred to *S. aureus* 6850 by phage transduction.

### RNAseq

RNA was isolated from *S. aureus* using TriZOL as previously described (21). DNase I treatment was performed to remove remaining DNA. The concentration of the RNA was determined by spectrophotometry on a Nanodrop 1000 (Peqlab) and RNA integrity was examined using agarose gel containing formamide or by analysis via Bioanalyzer. Enrichment of mRNA was done using the Universal Ribodepletion Kit followed by Next Ultra Directional Library Preparation Kit for Illumina (NEB). The cDNA was sequenced on HiSeq™ 2500 (Illumina) yielding 100 bp paired end reads. Adapters were removed using cutadapt (22) and only reads exceeding a mean base quality 5 within all sliding windows of 5 bp were mapped to the *S. aureus* 6850 genome (NCBI accession NC_022222; (23)). Read mapping was conducted using Bowtie2 (24). DeSEQ2 (18) was used to identify differentially regulated transcripts.

### qRT-PCR

Reverse transcription of total isolated RNA was performed using RevertAID reverse transcriptase (Thermo Scientific). A 10-ng sample of cDNA was used to perform qRT-PCR in a one-step reaction using Sybr Green master mix (Genaxxon) on a StepOne Plus real-time PCR system (Applied Biosystems). For primers used for qRT-PCR see Table S2. Analysis was performed using the 2^−ΔΔ*CT*^ method. Relative gene expression was normalized to expression of *gyr*B and to the corresponding expression in control conditions.

### Statistics

Data were analyzed by Microsoft Excel and GraphPad Prism. Significance of differences of at least three independent replicates were determined by a Student’s t-test for comparison of two independent data sets or with one-way ANOVA followed by Tukey’s multiple comparison test for comparison of three independent data sets.

## Results

### TnSeq identifies a novel transcriptional regulator of *S. aureus* required for colonization of secondary organs

In order to identify factors required for *S. aureus* colonization of secondary organs during bloodstream infection, mice were intravenously infected with 10^6^ CFU of a transposon mutant library of *S. aureus* strain 6850 (11), liver and kidneys were isolated from infected mice (n=3) at 24 h after bacterial inoculation and organ homogenates were plated onto blood agar. The number of bacteria at the time of sampling was 4.86±0.69 Log_10_ CFU in liver and 5.3±0.7 Log_10_ CFU in kidneys. Bacteria were collected from the plates, pooled, and enrichment or depletion of bacterial insertional mutants in kidneys and livers of infected mice was assessed by deep-sequencing transposon insertions sites (Tn-Seq).

Of five genes (RSAU_000958, *pur*M; RSAU_000901, RSAU_000571, a hypothetical protein RSAU_000852, and RSAU_002542, a putative DNA-binding protein) whose mutants were significantly depleted in bacteria recovered from kidneys in comparison with the inoculated mutantpool (Table 1) four were also found depleted in liver: RSAU_000958, RSAU_000901, RSAU_000571, and RSAU_002542 (Table S4, Table S5, Table S6). Whereas *pur*M and ItaA are genes with known functions and involved in pathways that are important for the physiology of *S. aureus* such as the purine biosynthesis and cell wall, respectively, RSAU_000852 (NCTC8325 Identifier SAOUHSC_00913) was a novel gene with unknown function. Because this gene is homologous to a gene encoding a LysR-type transcriptional regulator we termed RSAU_000852 as LysR-type transcriptional regulator (LTTR). Mutants in LTTR were significantly depleted in bacteria recovered from kidneys (log_2_FC 3.95, p=5.6 × 10^−7^), whereas read counts obtained from liver did not differ significantly from those of the inoculum (log_2_FC=0.18, p=0.84) (Fig. 1). Sequence alignments indicated that LTTR is highly conserved in *S. aureus*, where sequence variants share at least 96% identity on the amino acid level. In addition to *S. aureus*, homologues were also observed in the closely related species *S. schweitzeri* and *S. argenteus*, with slightly different gene variants being observed in *S. argenteus* (84-87% of amino acid sequence identity between *S. argenteus* and *S. aureus* proteins; data not shown). Since the gene is conserved in *S. aureus*, we focused our efforts on the functional characterization of this novel regulator that seems to be critical for the fitness of *S. aureus* for the colonization of secondary organs. For this purpose, we generated an isogenic knock-out mutant within the *S. aureus* strain 6850 deficient in the expression of LTTR (Δ852). By genome-sequencing we excluded the introduction of secondary site mutations during the mutagenesis procedure, since the method involved selection of an integrative plasmid under non-permissive conditions (treatment at 42 °C), which had been shown to result in mutations within the sae two component system.

**Table 1.** List of genes identified based on the enrichment (positive log_2_-fold changes [log_2_FC] or depletion (negative log_2_FC of the respective transposon mutant in murine kidneys respect to the intravenously inoculated transposon mutant pool (cutoff: adj. p-Values <= 0.001.

| Locus ID | log<sub>2</sub>FC | Adjusted p-Value | gene/annotation |
| --- | --- | --- | --- |
| RSAU_000958 | -5.00 | 1.50E-04 | purM |
| RSAU_000901 | -4.11 | 1.24E-04 | ltaA |
| RSAU_000571 | -4.04 | 1.24E-04 | hypothetical protein |
| **RSAU_000852** | **-3.95** | **5.60E-07** | **LTTR** |
| RSAU_002542 | -2.94 | 1.24E-04 | DNA-binding protein, putative |
| RSAU_000494 | 3.72 | 2.75E-04 | rpoB |
| RSAU_000940 | 3.74 | 2.40E-04 | atl |
| RSAU_000222 | 3.96 | 1.24E-04 | lytM |
| RSAU_000981 | 4.09 | 2.85E-04 | pdhD |
| RSAU_001882 | 4.26 | 3.88E-04 | Gcp |
| RSAU_000151 | 4.33 | 4.86E-04 | Putative transporter, permease component transport system permease protein |
| RSAU_001970 | 4.40 | 2.62E-04 | ATP-grasp domain protein |
| RSAU_002089 | 4.43 | 2.02E-04 | pbuG |
| RSAU_000495 | 4.50 | 7.74E-04 | rpoC |
| RSAU_001555 | 4.51 | 4.52E-04 | pyk |
| RSAU_000943 | 4.53 | 6.32E-04 | nanE |
| RSAU_002274 | 4.53 | 1.89E-04 | bar |
| RSAU_000822 | 4.57 | 2.91E-04 | Pyridine nucleotide-disulfide oxidoreductase family protein |
| RSAU_001560 | 4.81 | 1.24E-04 | dnaE |
| RSAU_000844 | 4.84 | 1.24E-04 | addA |
| RSAU_002334 | 5.00 | 7.83E-04 | pgcA |
| RSAU_000006 | 5.04 | 1.76E-06 | gyrA |
| RSAU_000735 | 5.04 | 8.10E-04 | hprK |
| RSAU_000187 | 5.08 | 2.36E-04 | ptsG |
| RSAU_000193 | 5.10 | 2.36E-04 | gutB |
| RSAU_001351 | 5.14 | 1.24E-04 | ebpS |
| RSAU_002484 | 5.15 | 5.55E-04 | manP |
| RSAU_002525 | 5.16 | 3.05E-04 | hisZ |
| RSAU_000259 | 5.17 | 9.20E-04 | nupC |
| RSAU_000009 | 5.19 | 1.24E-04 | serS |
| RSAU_000666 | 5.22 | 1.50E-04 | Transporter Anion:Sodium Symporter family protein |
| RSAU_002543 | 5.25 | 1.24E-04 | nixA |
| RSAU_001219 | 5.28 | 2.50E-04 | guaC |
| RSAU_000600 | 5.31 | 7.74E-04 | nupC2 |
| RSAU_002444 | 5.35 | 1.17E-04 | Amino acid permease family protein |
| RSAU_000994 | 5.58 | 4.08E-07 | typA |
| RSAU_000042 | 5.59 | 2.28E-04 | Dienelactone hydrolase family protein |
| RSAU_002289 | 5.60 | 1.96E-04 | rpsP |
| RSAU_001080 | 5.62 | 7.42E-04 | rluD |
| RSAU_002278 | 5.70 | 4.86E-04 | cycA |
| RSAU_000062 | 5.76 | 3.24E-05 | 67 kDa myosin-cross-reactive antigen |
| RSAU_001145 | 5.77 | 2.91E-04 | proS |
| RSAU_001630 | 5.84 | 7.83E-04 | Mannosyl-glycoprotein endo-beta-N-acetylglucosaminidase protein |
| RSAU_000638 | 5.94 | 1.24E-04 | vraG |
| RSAU_000864 | 5.97 | 4.22E-04 | oppA |
| RSAU_000268 | 5.98 | 6.89E-04 | yqiG |
| RSAU_002020 | 6.21 | 2.02E-04 | htsB |
| RSAU_000696 | 6.23 | 5.21E-04 | ptsH |
| RSAU_000631 | 6.32 | 1.24E-04 | lipA |
| RSAU_000427 | 6.50 | 2.40E-04 | metG |
| RSAU_002217 | 6.51 | 9.28E-09 | rsp |
| RSAU_002432 | 6.60 | 3.09E-04 | Adenine nucleotide alpha hydrolases superfamily protein |
| RSAU_001217 | 6.78 | 9.15E-04 | rpmG |
| RSAU_002292 | 7.11 | 3.22E-07 | rimM |
| RSAU_000505 | 7.24 | 6.55E-05 | capD |

**Fig. 1:**
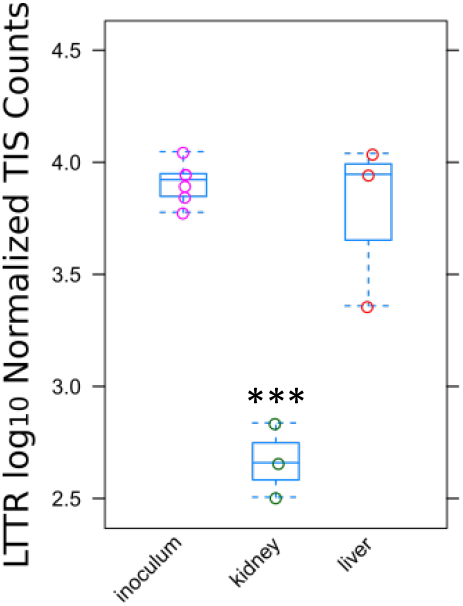
The frequency of *S. aureus* with mutation in RSAU_000852 is significantly lower in the bacterial pool recovered from infected kidneys than in the original inoculum. C57BL/6 mice (n=3) were intravenously infected with a *S. aureus* 6850 transposon mutant library and viable bacteria were recovered from liver and kidneys at 24 h of infection. Pools of the recovered bacteria and the respective inoculum were analyzed by Tn-Seq and counts of transposon insertions sites were compared by DEseq2. ***, *p* < 0.001.

### LTTR is important for *S. aureus* successful colonization of secondary infection sites

To validate the results of the transposon mutant screen, we compared the capacity of *S. aureus* Δ852 to establish infection in secondary organs after intravenous inoculation with that of the wildtype strain. Whereas CFUs of wildtype and mutant strains recovered from liver did not differ significantly (Fig. 2A), the numbers of bacteria were recovered from kidneys (Fig. 2B) and bones (Fig. 2C) were significantly lower in mice infected with Δ852 than in those infected with the wildtype strain. The serum levels of the inflammatory cytokine IL-6 were significantly lower in mice infected with Δ852 than in mice infected with the wildtype strain, further corroborating the attenuated phenotype of Δ852 (Fig. 2D). These results supported the requirement of LTTR for *S. aureus* successful infection of secondary organs during bloodstream infection.

**Fig. 2:**
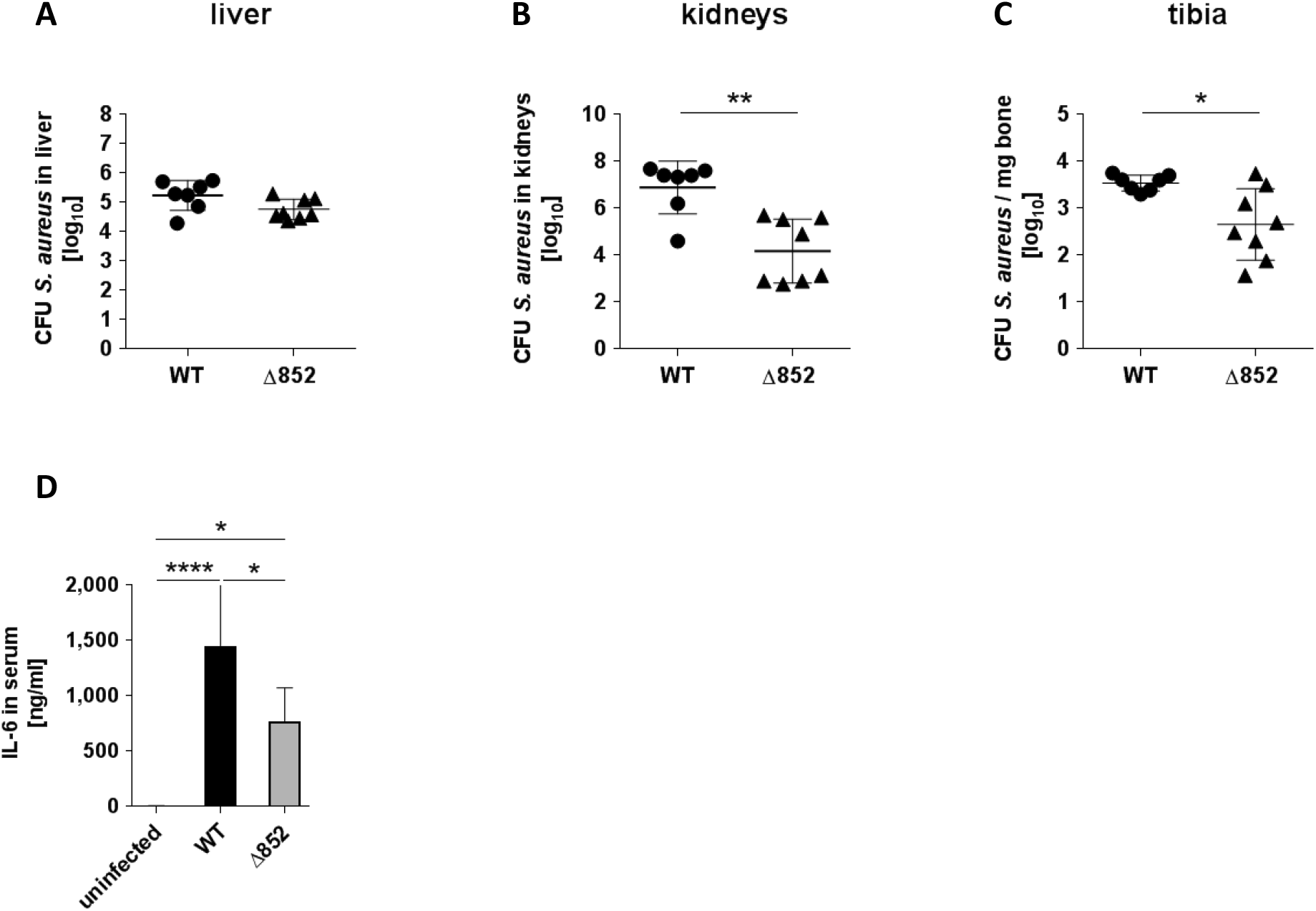
LTTR is required by *S. aureus* for successful infection of secondary organs. Mice were infected intravenously with wildtype (circle) and Δ852 (triangles) and CFU where determined in (**A**) liver, (**B**) kidneys and (**C**) tibia at 24 h of infection. Each symbol represents the data of an individual animal. Data are the pool of three independent experiments. (**D**) Serum levels of IL-6 in uninfected or in mice infected with either wildtype or Δ852 mutant strain at 24 h of infection. Each column represents the mean values ± SD of three independent experiments. *, *p* < 0.05; **, *p* < 0.01; ****, *p* < 0.001.

### Characterization of the regulon of the novel LTTR

To identify the regulon controlled by LTTR, *S. aureus* wildtype (WT) and *S. aureus* Δ852 were transduced with an anhydrous tetracycline (AHT)-inducible plasmid (25) carrying LTTR (WT_AHT and Δ852_AHT, respectively). The gene expression profile of *S. aureus* WT, Δ852, WT_AHT, Δ852_AHT cultured in TSB medium and was analyzed using RNA-seq after inducing LTTR expression by addition of 200 ng/ml of AHT for 1 h and the genes differentially expressed among the different strains were assessed by DEseq2 analysis. Several transcripts were found specifically up- and downregulated upon induction of LTTR (Table 2). Genes, upregulated upon induction of LTTR expression included peptide methionine sulfoxide reductase *msr*A2, the components of a copper-translocation machinery *cop*A and *cop*Z as well as the genes of branched chain amino acid biosynthesis *ilv*D, *ilv*B, *leu*A2 and the holin-like murein hydrolase regulator *lrg*A. The genes with reduced expression after LTTR induction included members of the urease operon (*ure*B, *ure*C, *ure*E and *ure*F), genes of pyrimidine biosynthesis pathway (*pyr*B, *pyr*C and *car*B), the gene coding for Tagatose-6-phosphate kinase (*lac*C) and PTS system lactose-specific transporter subunit IIBC (*lac*E). The upregulation of LTTR in Δ852_AHT after treatment with AHT was validated by RT-PCR (Fig. 3A). Furthermore, the expression of a set of genes found in the RNA-seq data either upregulated (*msr*A, *cop*A, *lrg*A and RNAIII) or downregulated (*leu*A2, *ure*C, *leu*A1, *car*B, *lac*C and *pyr*B) after LTTR induction in WT-AHT versus Δ852 were also confirmed by RT-PCR (Fig. 3B).

**Table 2:**
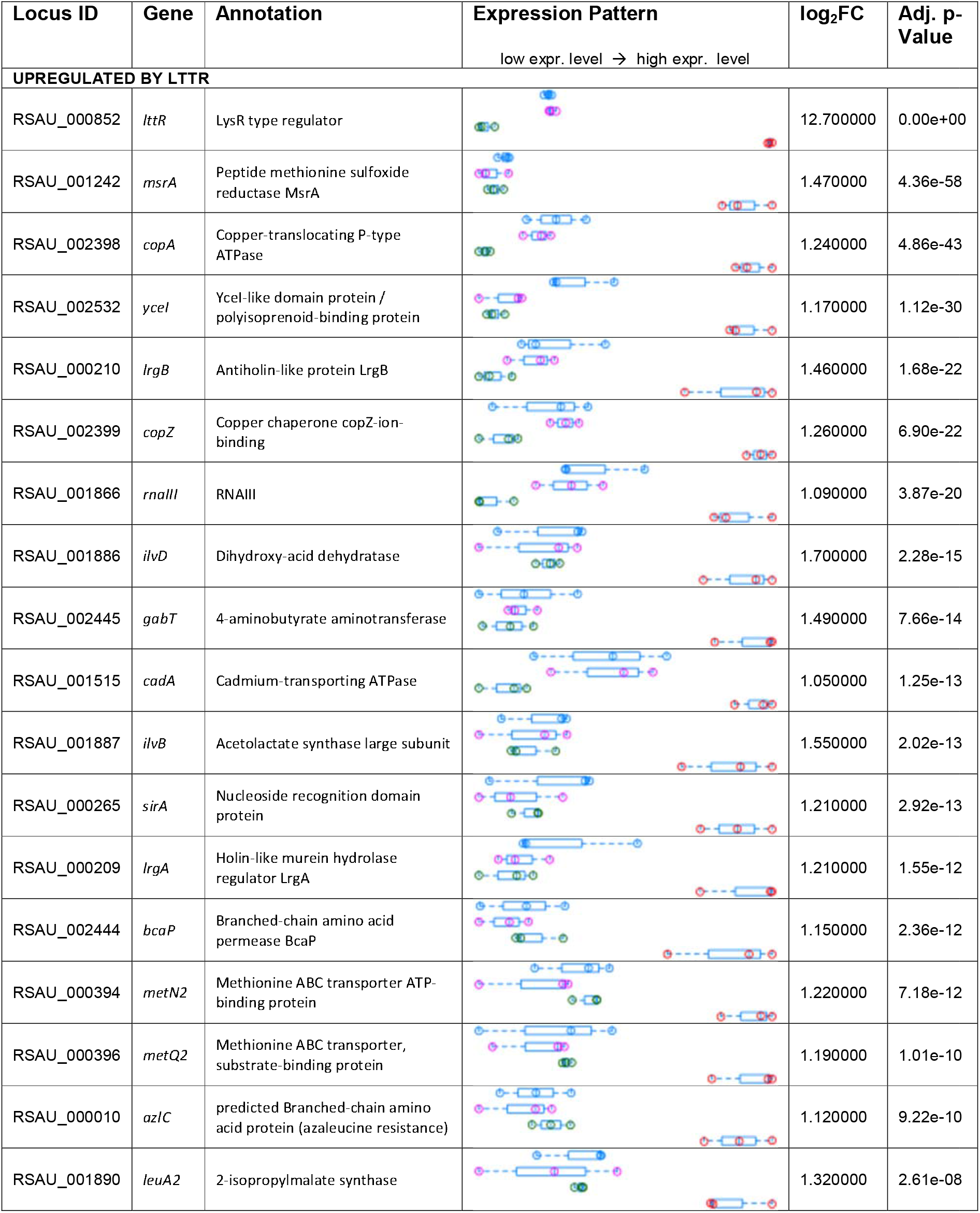
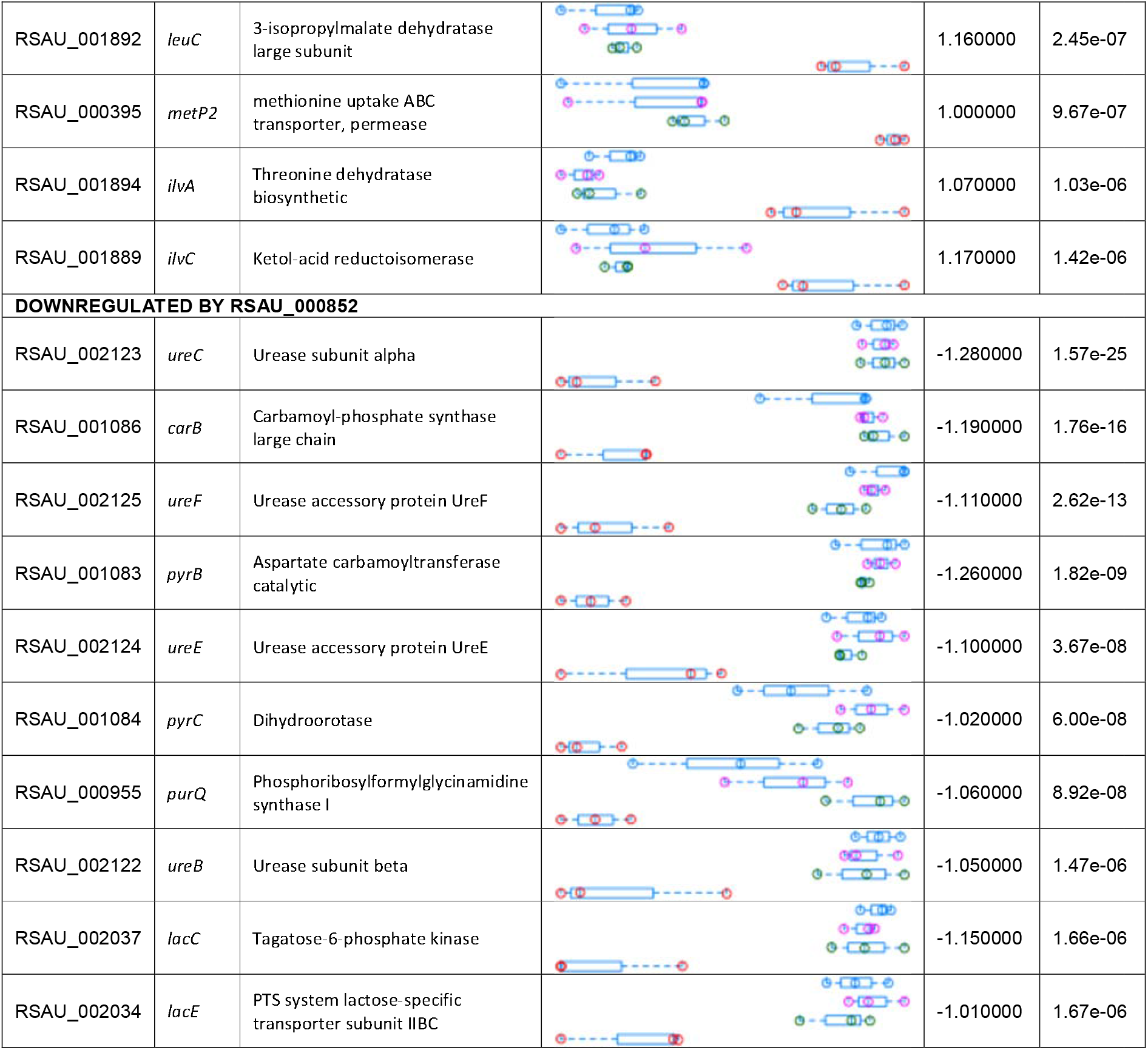
Regulated genes upon induced expression of LTTR. The Expression pattern is color coded: Blue: *S. aureus* WT, Magenta: WT treated with 200 ng/ml AHT, Green: *S. aureus* Δ852, Red: *S. aureus* pI852, LTTR expression induced with 200 ng/ml AHT.

**Fig. 3:**
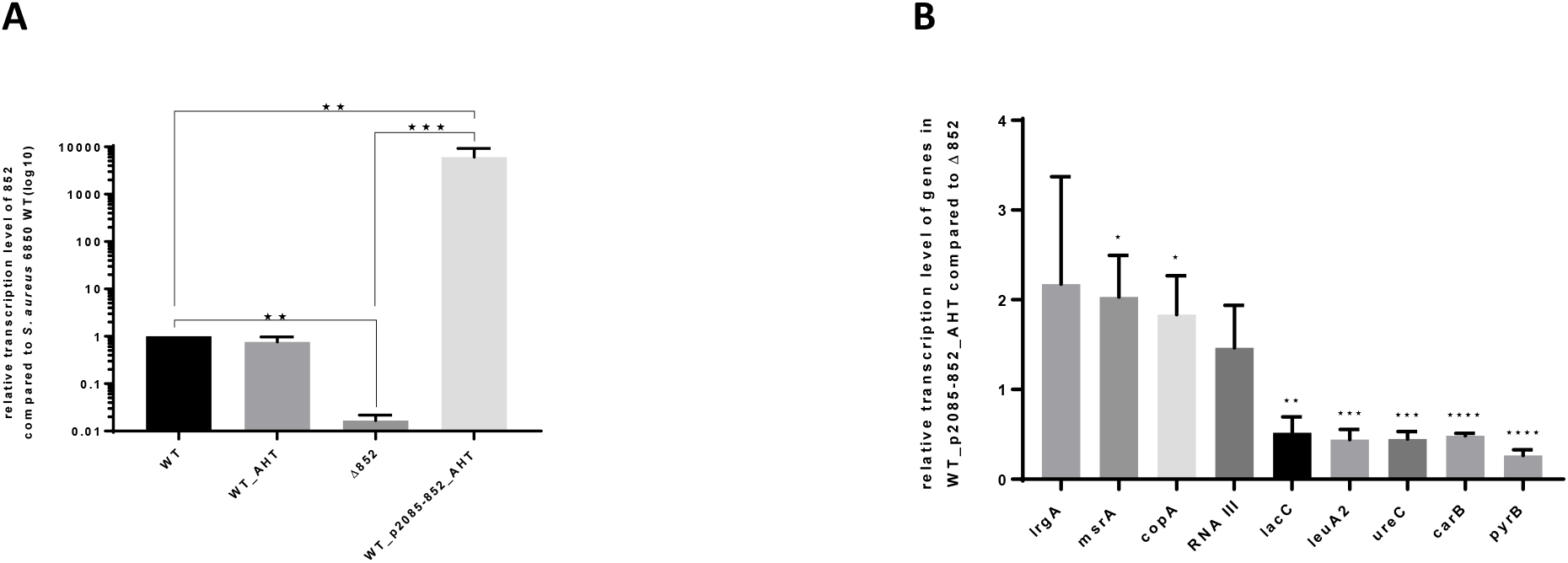
Validation of RNA-seq data by quantitative RT-PCR. (**A**) Expression level of LTTR (RSAU_000852) in *S. aureus* Δ852, WT_AHT and Δ852_AHT relative to the expression in the WT strain after stimulation of LTTR expression with 200 ng/ml AHT for 1 h (n=3). Notice the similar levels of LTTR expressions in WT and WT_AHT and the almost undetectable levels in Δ852. (**B**) Expression level of a set of genes in WT_AHT relative to the expression in Δ852. Statistical analysis was performed using Student’s t-test. *, *p* < 0.05; **, *p* < 0.01;***, *p* < 0.001;****, *p* < 0.0001.

Using the genes’ ortholog identifiers for strain NCTC8325, we found that the potential LTTR target genes were significantly enriched in GO-terms for the biological processes “branched-chain amino acid biosynthetic process” (GO:0009082), “urea metabolic process (GO:0019627) and ‘de novo’ UMP biosynthetic process (GO:0044205) (Table S3).

### Transcription of LTTR is enhanced under infection-mimicking conditions

To investigate under which conditions LTTR is expressed by *S. aureus*, a promoter fusion with a codon-adapted green fluorescent protein (GFP) (26) was generated yielding the plasmid p2085-Pr852-GFP that was introduced into *S. aureus* WT. Because oxygen levels at sites of tissue inflammation may be relatively low as a result of impaired perfusion (27), we recorded bacterial growth and GFP fluorescence levels under aerobic and microaerobic conditions. No differences were observed in bacterial growth under aerobic or microaerobic conditions (Fig. S1A). However, promoter activity was found stronger in bacteria grown under microaerobic conditions when compared to aerobic conditions (Fig. S1B, C). We therefore determined the levels of transcription of LTTR in *S. aureus* WT grown in TSB under aerobic and microaerobic conditions and found the transcription levels of the gene to be significantly higher under microaerobic conditions than when grown in presence of oxygen (Fig. S1D).

Interestingly, during the process of generating the GFP reporter strain, we observed that *S. aureus* Pr852-GFP plated on LB agar without glucose exhibited more intense GFP fluorescence than when it was grown in TSB agar containing 2.5 g/L glucose (Fig. 4A). Similarly, we observed difference during growth in broth as shown by the slightly faster bacterial growth in glucose-free LB-Medium when compared to growth in TSB-Medium containing 2.5 g/L glucose. The mid-log phase was reached after 3 h in LB and after 7 h culture in TSB (Fig. S1B). Again, promoter activity of Pr852 was drastically increased in LB in comparison to TSB (Fig. S1C, D). In order to investigate the repression by glucose, we grew WT and mutant bacteria in chemically defined medium (CDM) containing various concentrations of glucose ranging from 0 to 75 mM. We recorded bacterial growth as well as Pr852 promoter activity over 24 h (Fig. S1E). Addition of 10 mM glucose resulted in ~50% reduction of promoter activity, which was almost completely abolished at 50 mM and 75 mM glucose. Our results hereby indicate that the promotor of LTTR is repressed by glucose.

**Fig. 4:**
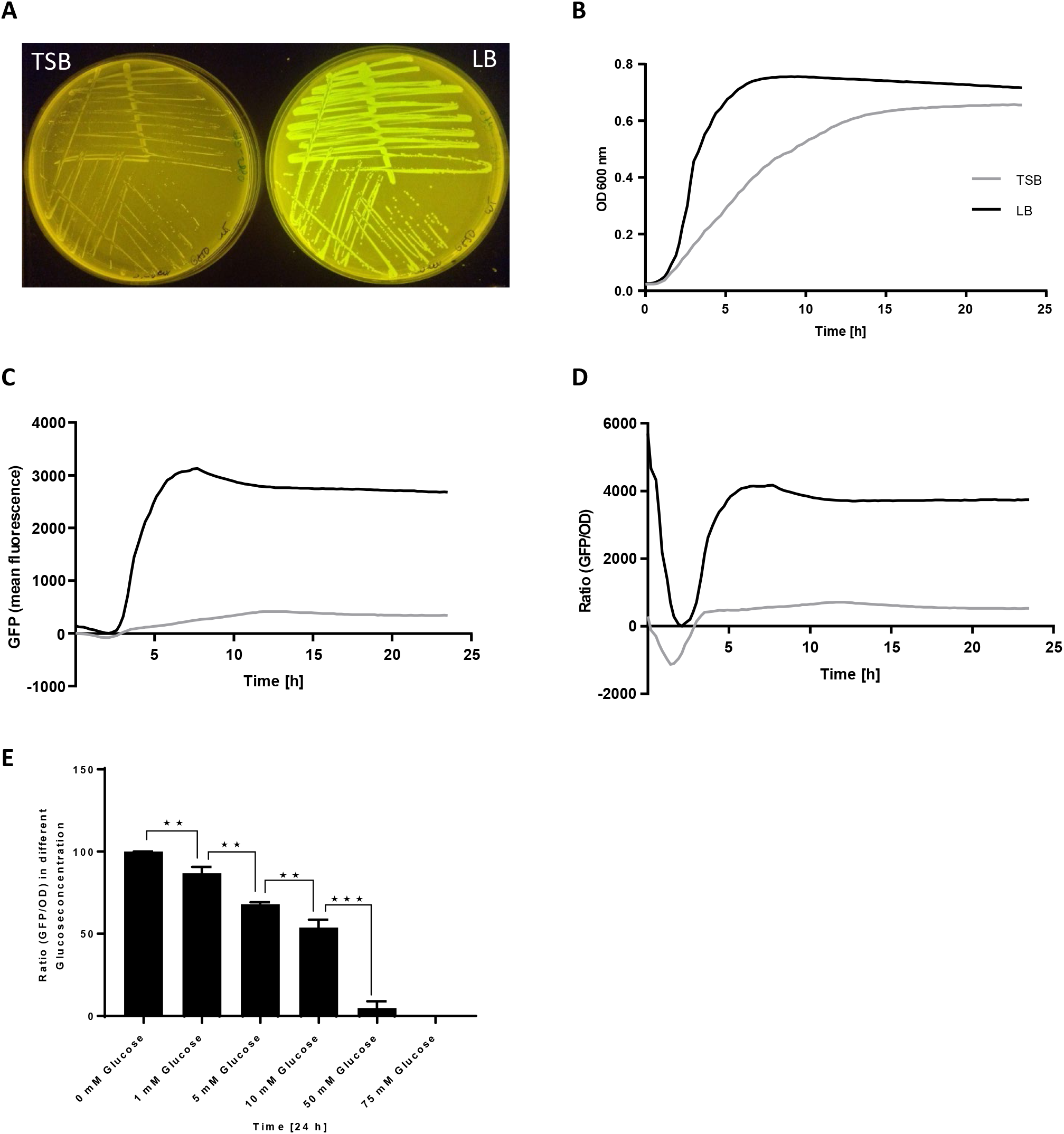
Transcription of LTTR is repressed by glucose. (**A**) Green GFP fluorescence of *S. aureus* 6850 harboring the LTTR promoter reporter plasmid Pr852-GFP is less intense in on TSB-agar containing 2.5 g/L glucose (left) when compared to glucose-free LB-Agar (right). (**B-D**) Kinetics of *S. aureus* 6850 Pr852-GFP growth determined over 24 hours of growth under microaerobic conditions in either glucose-containing TSB or glucose-free LB-Medium. (**B**) OD600, (**C**) Mean GFP fluorescence. (**D**) Ratio GFP/OD. (**E**) GFP/OD ratios after 24 h of culture of the promoter reporter strain grown under microaerobic conditions in CDM supplemented with different glucose concentrations demonstrate a concentration-dependent repression of the LTTR promoter by glucose. Each kinetics experiment was performed in triplicate (n=3). Statistical analysis was performed using Student’s t-test. **, *p* < 0.01;***, *p* < 0.001

In order to investigate additional nutritional requirements for LTTR transcription, we used CDM and determined LTTR promoter activity by the levels of GFP fluorescence. We first corroborated that promoter activity was enhanced under microaerobic conditions in CDM medium (Fig. S1E) and increased in the absence of glucose (Fig. 5A). We next determined LTTR promoter activity in CDM without glucose and omitting other components of the defined medium. Upon omission of the trace metal mixture, we identified a reduced LTTR activity (Fig. 5A). We thus omitted single constituents of the trace mixture including boric acid, manganese (II) chloride, zinc sulfate, sodium molybdat, cobalt nitrate, or copper sulfate (Fig. 5B, S2A-F). LTTR promoter activity was abolished only in CDM without copper sulfate, but showed a dose-dependent increase upon addition of copper sulfate which was fully reconstituted by addition of at least 400 nM copper sulfate (Fig. 5C, D). Similarly, the relative transcription level of LTTR in strains grown in CDM without glucose (Δglucose) compared to CDM additionally lacking copper sulfate (CDM Δglucose Δcopper) demonstrated that the transcription level of LTTR is significantly higher in *S. aureus* WT (ΔΔ^CT^=2.375± 0.1737, p-value=0.0014) as well as Δ852 bacteria carrying the complementation plasmid pI852 (ΔΔ^CT^=2.527 ± 0.1795, p-value=0.0010) when copper sulfate was present in the media (Fig. S2GG).

**Fig. 5:**
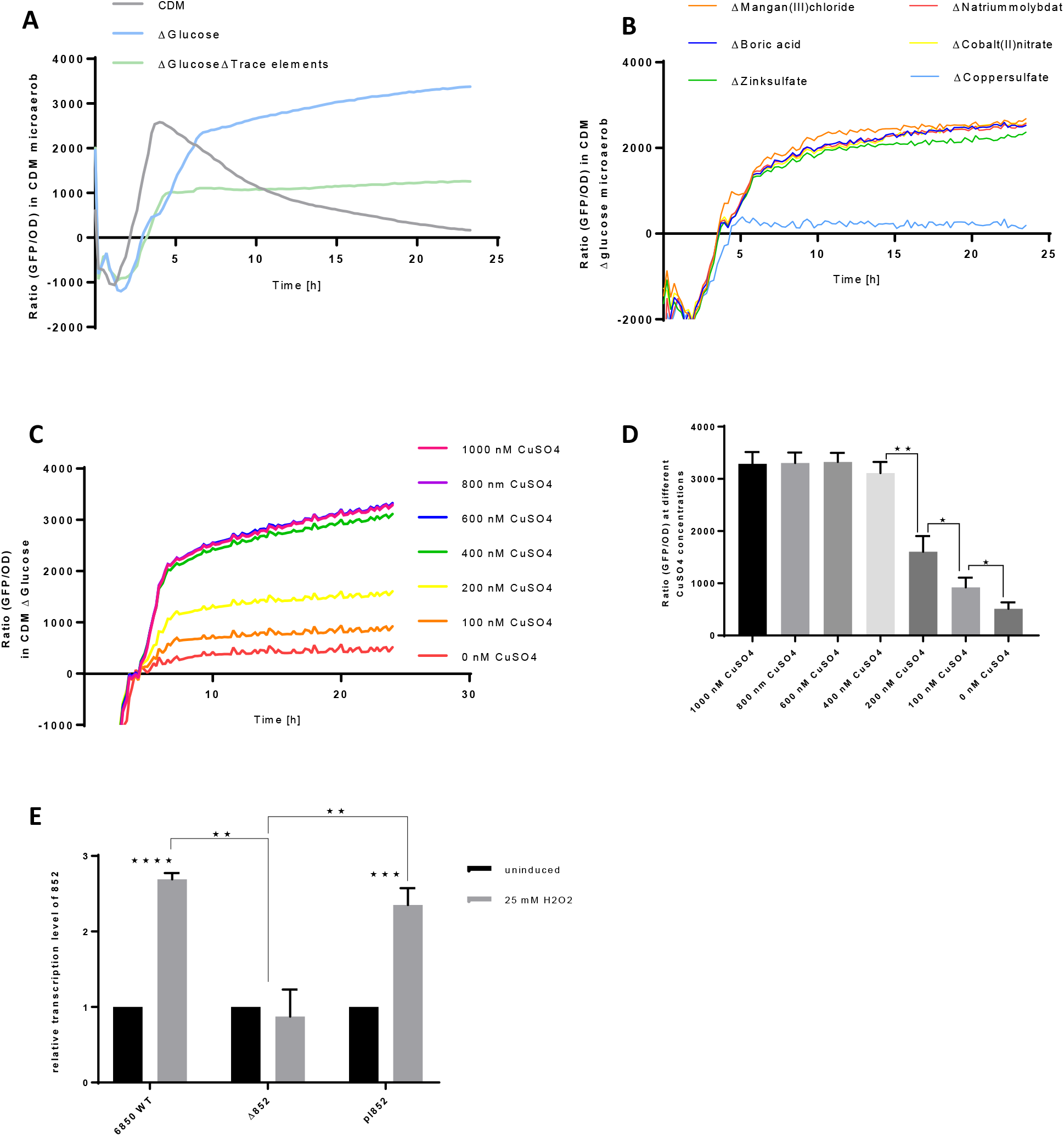
LTTR promoter activity is observed under infection-mimicking conditions and requires copper ions. (**A**) GFP / OD ratios of wildtype *S. aureus* 6850 growing under microaerobic conditions in full CDM (light grey), CDM lacking glucose (blue), or lacking both, glucose and trace elements (green). (**B**) Microaerobic growth in glucose-free CDM lacking single constituents of the trace element mixture identifies a copper requirement for promoter activity. (**C**) Copper-dependent dose response of the LTTR promoter is observed during microaerobic growth in glucose-free CDM and increasing amounts of copper sulfate ranging from 0-1000 nM. (**D**) 400 nM copper sulfate restore full promoter activity, whereas lower concentrations show dose-dependent significant decrease in promoter activities. (**E**) The LTTR promoter is induced by 25 mM hydrogen peroxide. Relative transcription level of the different strains under un-induced (black bar) conditions or after treatment with 25 mM H_2_O_2_ for 1h (grey bar). Each experiment was performed in triplicates (n=3). Statistical analysis was performed using Student’s t-test. *, *p* < 0.05; **, *p* < 0.01;***, *p* < 0.001;****, *p* < 0.0001.

Our RNAseq data after induced expression of LTTR identified the oxidative stress reponse gene *msr*A2 as a target gene. We therefore hypothesized that LTTR may be important for *S. aureus* upon encounter of oxidative stress in the host. To test this hypothesis, we compared the level of LTTR gene RSAU_000852 transcription between *S. aureus* WT, the Δ852 mutant as well as the complemented mutant grown under normal growth conditions and under oxidative stress conditions generated after a pulse with 25 mM hydrogen peroxide for 1 h. Transcriptional expression of the well-known oxidative stress response genes *dps* and *sod*A was used to corroborate that peroxide exposure induced an oxidative stress response in *S. aureus* (Fig. S3). Transcription of RSAU_000852 was induced in WT (2.69±0.05-fold induction; p-value<0.0001) and complementant (2.35±0.13-fold induction; p-value=0.0004) under H_2_O_2_ stress, which is absent in the Δ852 mutant (Fig. 5E).

## Discussion

After invasion of the host, pathogens needs to rapidly adjust to the different tissue microenvironments encountered during the infection process in order to successfully establish infections. This involves sensing local conditions, including nutrient availability and resident host immune defenses, an adjust their gene expression and virulence functions accordingly. Thus, within the infection niches encountered during infection, pathogens needs to overcome barriers that sharply limit the population size, the so-called bottlenecks. For example, after intravenous inoculation, *S. aureus* quickly accumulate within the liver, which constitutes the first infection niche of the host. Within the liver, a high proportion of the inoculated bacteria are sequestered by Kupffer cells, which are liver macrophages and represent the first infection bottleneck. Although a proportion of the bacteria sequestered within Kupffer cells are efficiently killed, a sub-population of internalized *S. aureus* bacteria have been shown to survive and even multiply in the phagosomes and eventually kill the host cells (28). There, neutrophils take up the bacteria and may reenter the circulation thereby disseminating the pathogen from the liver to secondary infection sites (29). From the first niche in the liver, *S. aureus* disseminates to secondary infection niches such as the bones and kidneys, where distal abscesses are formed (6). It is thought that *S. aureus* requires different subsets of virulence factors to establish infection in the various niches it may encounter in the host.

### TnSeq identified a niche specific transcription factor

We hence used a transposon mutant library in *S. aureus* 6850 (30) to discover virulence factors critical for *S. aureus* to establish infection in secondary infection niches. Mice were intravenously inoculated with the transposon library and Tn-seq was applied to bacteria recovered from the primary infection site at the liver as well as from secondary infections site at the kidneys to catalog the insertion mutants when compared to the infection inoculum. Several genes were identified whose mutants were specifically depleted or enriched in either liver or kidney or in both organs of mice after experimental intravenous infection (Table 1, S5, S6).

Interestingly, the lists obtained by Tn insertion site sequencing do not contain classical (pore-forming) toxins such as staphylococcal α-toxin or leukocidins, illustrating that the sequencing strategy would benefit from mutant libraries with higher complexities (12), however, the strategy is mostly limited by the strong selective effects. For instance, we initially tried to extend the TnSeq strategy to tibiae, however, the recovered number of bacteria was too low for analysis thereby illustrating the strength of the bottleneck effects (data not shown).

We found only the RSAU_000852 mutant to be specifically depleted in secondary infection sites in kidneys in comparison with the original inoculated mutants pool as well as with those mutants recovered from primary infection sites at the liver. These results indicate that expression of RSAU_000852 may be of critical importance for *S. aureus* to establish infection specifically in secondary niches within the host. This was corroborated by intravenous infection of mice with the seamless deletion mutant Δ852 deficient in the expression of RSAU_000852. Whereas Δ852 mutant exhibited significantly lower capacity to establish infection in kidneys and tibiae than wild type bacteria, the number of Δ852 mutant bacteria recovered from liver was comparable to that of wild type strain. Furthermore, mice infected with the Δ852 mutant also exhibited significantly lower serum levels of the inflammatory cytokine IL-6 than those infected with the wild type strain, which correlates with the attenuated phenotype of the mutant strain.

### Characterization of the novel LTTR

Homology research identified a LysR-type transcriptional regulator (LTTR) as the product of RSAU_000852. LTTRs are known as the largest family of prokaryotic DNA-binding proteins, with 800 members identified on the basis of their amino acid sequence (31). They can act as either repressors or activators of single or operonic genes (32, 33). LTTRs regulate numerous genes, whose products are involved in important bacterial functions such as metabolism, motility, cell division, quorum-sensing, virulence, oxidative stress response, to name a few (34–42). Four LTTRs have been identified so far in *S. aureus* including CidR, HutR, GltC and CcpE (43–45). In this study, we have identified a new LTTR in *S. aureus* 6850 encoded by RSAU_000852, with unknown function and whose expression was rarely reported. For example, some studies have reported that RSAU_000852 was 2.94-fold downregulated in presence of 0.01 μM linoleic acid (46) and 3.75-fold induced in strain COL exposed to 1 mM diamide (47). In other studies, however, where networks of gene regulation were investigated in *S. aureus* strain HG001 under 106 different conditions, the NCTC82325 LTTR_852 homolog, SAOUHSC_0913, was never found to be expressed in any of the conditions tested (48).

By using AHT-inducible expression of LTTR in *S. aureus* 6850 and subsequent RNA-seq, we established potential targets of the transcriptional regulator (Table 2). Genes found upregulated upon induction of LTTR expression included those encoding peptide methionine sulfoxide reductase *msr*A2, components of a copper-translocation machinery *cop*A and the copper chaperone *cop*Z (49), which are involved in copper efflux, as well as genes encoding factors involved in the branched chain amino acid biosynthesis such as *ilv*D (Dihydroxy-acid dehydratase), *ilv*B, *leu*A2 and the holin-like murein hydrolase regulator *lrg*A. Oxidation of methionine by the activity of methionine sulfoxide reductase can destroy protein structure or affect protein function (50). There are four methionine sulfoxide reductases encoded by *S. aureus* of which three possess MsrA-functionality, and a single gene encoding MsrB (51). Interestingly, the operon encoding *msr*A1 and *msr*B was seen upregulated when comparing the transcriptomes of *S. aureus* grown in medium *in vitro* with *S. aureus* recovered from experimental animals during acute and chronic infection (6). Our data suggest, however, that LTTR is specifically upregulating *msr*A2. Among the well-studied LTTRs is OxyR, which is responsible for the oxidative stress response against H_2_O_2_ in *Escherichia coli*, *Salmonella enterica* servoar Typhimurium (52). Peroxide is, for instance, produced as antibacterial agent by many cells of the innate immune system. Hence LTTR may be important for *S. aureus* upon encounter of oxidative stress in the host.

Isopropylmalate synthase 2 encoded by *leu*A2 is involved in the first step of the pathway of L-leucine biosynthesis. *leu*A2 as well as other genes of the BCAA biosynthesis machinery (*leu*C, *leu*D, *leu*B and *ilv*E) and *bca*P, a BCAA transporter (53), have been identified as target genes of LTTR by RNA-seq. Another virulence-associated BCAA transporter, brnQ2 (RSAU_000252), was reported to be upregulated in *S. aureus* during chronic infections suggesting CodY derepression and emphasizing the importance of branched chain amino-acid metabolism for *S. aureus* survival within the host (6). *brn*Q1, *brn*Q2, or *brn*Q3 and other CodY targets, however, were not significantly regulated in our dataset, suggesting that a CodY-independent induction of BCAA biosynthesis may exist in *S. aureus*. It has to be noted that transcription of LTTR was not derepressed in a codY mutant (54), although the promoter region of LTTR (within the intergenic region of the upstream gene *clp*B and RSAU_000852) was highly enriched in CodY chromatin precipitation analyses (54).

Genes with reduced expression after LTTR induction included members of the urease operon *ure*B, *ure*C, *ure*E, and *ure*F as well as genes of pyrimidine biosynthesis *pyr*B, *pyr*C, and *car*B, suggesting that the *de novo* UMP biosynthesis pathway is repressed by LTTR. Since urease is part of the of *S. aureus* response to acid-shock and recently was shown to be required for bacterial persistence in murine kidney infection (55), it needs to be evaluated in which sub-organellar niche the wild-type and mutant bacteria are residing. *lac*C (Tagatose-6-phosphate kinase) and the lactose-specific compound *lac*E (PTS system lactose-specific transporter subunit IIBC) (56) were also found repressed by the LTTR. LacC is involved in synthesis of D-glyceraldehyde 3-phosphate and glycerone phosphate from D-tagatose 6-phosphate and LacE is part of the phosphoenolpyruvate-dependent sugar phosphotransferase system (PTS, LacEF). Increased PTS expression has been hypothesized to facilitate the pathogenesis of *S. aureus* in mastitis (57) and may also facilitate *S. aureus* adapatation to secondary organs during bloodstream infection as shown in this study.

Overall, our RNAseq expression data, although highly significant, demonstrated only moderate fold-change levels in the analysis. Since LTTRs require co-inducers for DNA-binding or multimerization, we assume that the concentration of the hitherto unknown co-inducer of LTTR was limiting under the conditions tested. LTTRs comprise an N-terminal helix–turn–helix (HTH) motif and a C-terminal LysR-substrate binding region (31). Different co-inducer with structural diversity (e.g. amino acid derivates, aromatics, ions, various aliphatics) have been shown to be involved in the stimulation of various LTTRs (31). For example, the well-characterized LTTR ToxR of *Burkholderia glumae* activates locally Quorum sensing via Toxoflavin (38). In rare cases, the co-inducer operates as an activator and repressor such as α-Methylene-γ-butyrolactone during pili and capsule synthesis in *Neisseria meningitidis* (58). Certainly, not all cofactors are known: for YofA involved in the cell division in *Bacillus subtilis* (41) and a *Yersinia pestis* co-repressor (59), the metabolites remain unidentified. Further experiments will be required to identify the co-inducer of LTTR. The decreased fitness of a RSAU_000852 mutant in mouse kidneys and tibiae leads us to hypothesize, that niche-specific small molecules may account for full activity of the LTTR protein.

Regarding the microenvironments in which LTTR is induced, our data show that LTTR promoter activity specifically requires microaerobic and glucose-free conditions, as well as low concentrations of copper ions. Kidneys are involved in glucose homeostasis through processes of gluconeogenesis, glucose filtration, reabsorption and consumption (60). Modeling glucose metabolism in kidneys identified anaerobic glucose metabolism in the inner medulla of glucose, owing to the limited blood flow and low tissue oxygen tension (61). Thus, microenvironments exist in the kidneys, which are compatible with promotor activity of RSAU_000852.

Furthermore, we identified copper as a limiting factor for the activation of LTTR transcription. Copper is a cofactor essential for many enzymes involved in a multitude of cellular functions of both host and pathogen, such as cellular respiration, iron transport, and free radical scavenging (62–64). Copper also serves as antibacterial agent and is pumped into phagosomes of macrophages in order to kill bacteria. *S. aureus*, in turn, possesses copper efflux pumps, which are involved in survival of high-copper stress. Recently, two novel genes were identified, copB and copL (65). *cop*B functions in copper export and displayed genetic synergy with *cop*A. The function of *cop*L is independent of *cop*A and *cop*B, it is a membrane bound and surface-exposed lipoprotein that binds up to four Cu-ions. Our data suggest that LTTR_852 not only is transcriptionally activated by low concentrations of copper ions, but also is involved in *cop*AZ expression thereby is modulating the copper response of *S. aureus*.

LTTR promoter activity can be seen in multiple genetic backgrounds of *S. aureus* (Fig. S4). Interestingly, copper depletion had no effect on the LTTR promoter activity in USA 300 JE2. USA 300 strains carry a copper resistance locus (copXL), uniquely associated with the SCCmec elements not found in other *S. aureus* lineages (66) (Fig. S4). *cop*L encodes a membrane-bound and surface-exposed lipoprotein that binds up to four Cu+ ions (65) and, therefore, may supply copper to the bacterium after been transferred to a copper-depleted growth media.

RNAseq after induced expression of LTTR suggested the oxidative stress reponse gene *msr*A as a target gene. We therefore hypothesized that LTTR may be important for *S. aureus* upon encounter of oxidative stress in the host (see above). Interestingly, we show induction of the LTTR gene RSAU_000852 under H2O2 stress, which is absent in the Δ852 mutant yet reinstated by supplying the locus in trans.

Taken together, our data show that the LTTR encoded by RSAU_000852 is a transcriptional regulator required in specific host niches for the efficient establishment of infection. Within this niche glucose and oxygen are limited, copper ions must be present, and reactive oxygen concentrations may be elevated, possibly due recruited immune cells.

Limitations of the study: We conducted TnSeq of *S. aureus* mutant pools recovered from liver and kidneys of intravenously infected animals. We identified the regulator LTTR as important for colonization of secondary organs (kidneys, bones) but not for establishment of infection in the primary site of infection at the liver. Since we harvested the infected tissues at a single time point very early during infection (24 h), we cannot discard a potential role of LTTR for bacterial survival in the liver at later times of infection.

Furthermore, expression analysis of potentially LTTR-regulated genes under induction conditionsd were not very high although significant. We hypothesize that the lack of the specific co-inducer that LysR-type transcriptional regulators ususally require for their activity is responsible for this effect.

In summary, we identified in this study a fifth LysR-type transcriptional regulator in *S. aureus* that is required for bacterial colonization of secondary infection sites. We unveilled a complex dependency of transcriptional activation of the regulator gene and by inducible expression identify a set of potentially regulated genes, which includes genes required to cope with oxidative stress and for growth in low-oxygen and glucose-free conditions as those found in host tissues. Hence, we hypothesize that the LTTR may play an important role in metabolic adaptation of *S. aureus* to local infection sites in the host. Thus, therapeutic targeting of the novel regulatory factor identified in this study in patients with *S. aureus* bacteremia may result in reduced bacterial fitness and offer options for improving disease outcome.

## Acknowledgements

We thank the German Research Foundation (Deutsche Forschungsgemeinschaft, DFG, www.dfg.de) for funding this work within the Collaborative Research Center TRR34 under code P11 to M.F. and T.R. and the University of Würzburg for financial support. We are indebted to Kerstin Paprotka and Linda Raupach for initial molecular cloning. This publication was funded by the University of Wuerzburg in the funding programme Open Access Publishing.

## Supplementary Materials

### Supplementary Tables

**Table S1: Bacterial strains and plasmids used in this study**

**Table S2: Oligonucleotides used in this study**

**Table S3: Enrichment in GO-terms as determined by STRING-DB.org**

**Table S4. Gene mutants identified enriched or depleted in murine liver after intravenous infection (cutoff: adj. p-Values <= 0.0005)**

**Table S5: *S. aureus* transposon mutants found either enriched or depleted in both, kidney or liver of mice after intravenous infection by TN-seq.** Counts of sequenced transposon insertion sites were compared with the counts obtained from the infection inoculum. By comparing this inocolum to the recovered samples genes which are lost in a specific tissue are represented by negative log_2_FC and vice versa.

**Table S6: *S. aureus* transposon mutants identified by Tn-seq with sequence reads enriched (positive log_2_-fold changes [log_2_FC]) or depleted (negative log_2_FC) in either kidney or liver of mice 24 hours after intravenous infection when compared to the infection inoculum (adjusted p-value cutoff 0.05)**. Shown are the ORF ID of strain 6850 as well as its homolog within strain NCTC8325, gene assignment, annotation, log2FC and adjusted p-value of reads obtained from bacteria recovered from infected kidneys and livers. Genes that were identified as infection-relevant in a previous study (67) are indicated.

### Supplementary Figure Legends

**Figure S1: Microaerophily has a positive effect on the promotor expression of LTTR**. (**A**) Kinetic of *S. aureus* WT growth determined by OD600 under microaerobic (lighter color) or under aerobic (darker color) conditions (n=3). (**B**) Mean fluorescence of the GFP linked to the LTTR promoter region in *S. aureus* WT under microaerobic (lighter color) or under aerobic (darker color) conditions. (**C**) Ratio of GFP to OD (GFP/OD) is higher under microaerobic conditions. (**D**) Relative transcription level of LTTR in *S. aureus* WT grown in TSB for 6 h under aerobic or microaerobic conditions. (**E**) GFP / OD ratios of wildtype *S. aureus* 6850 in CDM under microaerobic (light grey) or aerobic conditions (dark grey); Statistical analysis was performed using Student’s t-test. **, *p* < 0.01.

**Figure S2: LTTR852 Promotor activity under microaerobic conditions in CDM without glucose and trace elements illustrates the requirement of copper**. GFP / OD ratio of *S. aureus* p852 GFP in glucose-free CDM lacking either (**A**) boric acid, (**B**) manganese (II) chlorid, (**C**) zinc sulfate, (**D**) sodium molybdate, (**E**) cobalt (II) nitrate or (**F**) copper sulfate. (**G**) Significant higher expression of LTTR in wildtype and complementant is observed when strains are grown in CDM Δglucose in the presence of copper sulfate. Depicted are relative transcription levels of LTTR in wildtype (WT) and complementant (pl852) with or without copper sulfate. Mutant expression levels were set to one. Statistical analysis was performed using Student’s t-test. **, *p* < 0.01.

**Figure S3: LTTR transcription is induced by peroxide stress: Relative transcription levels of dps and sodA show induction of oxidative stress**. Relative transcription level of the *S. aureus* 6850 WT, the LTTR mutant (Δ852) and the complementation strain (pI852) under normal growth conditions (black bar) or after treatment with 25 mM H2O2 for 1h (grey bar). (A) General stress protein dps and (B) superoxide dismutase sodA are induced demonstrating a transcriptional response of *S. aureus* to reactive oxygen species. *, p < 0.05; **, p < 0.01;***, p < 0.001;****, p < 0.0001.

**Figure S4: LTTR promoter activity in different *S. aureus* strains in CDM under various conditions**. GFP/OD Ratios of (A) *S. aureus* 6850, (B) USA 300 JE2, (C) Cowan I and (D) and RN4220 in CDM under microaerobic conditions (black), CDM without glucose under microaerobic (green) and aerobic conditions (red), as well as in medium without glucose and copper sulfate (blue). All experiments were performed in triplicates (n=3).

