## Supplementary figures and images for "Identification of a novel LysR-type transcriptional regulator in *Staphylococcus aureus* that is crucial for secondary tissue colonization during metastatic bloodstream infection"

### Figure S1

#### Supplemental Figure 1

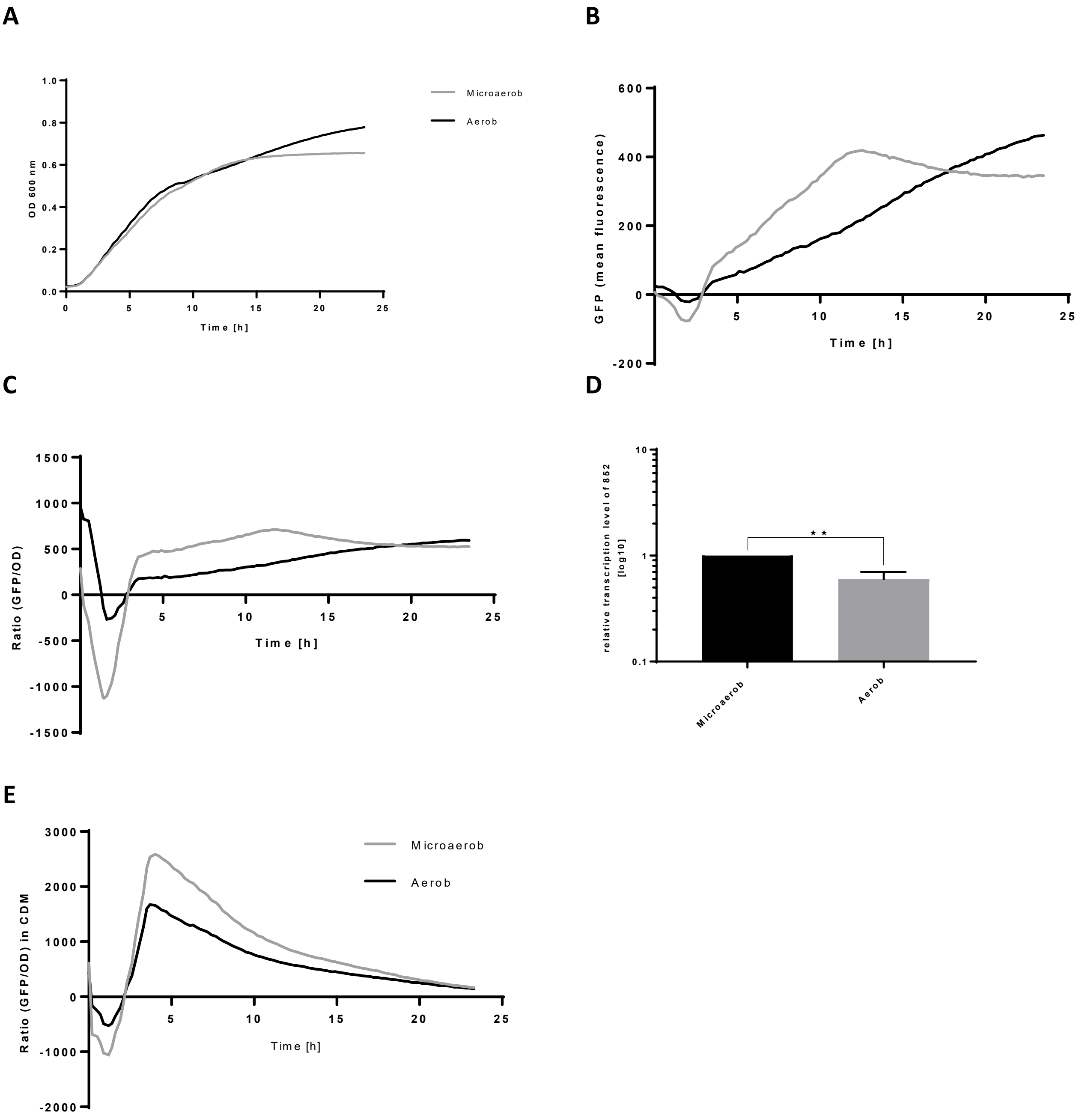

### Figure S2

#### Supplemental Figure 2

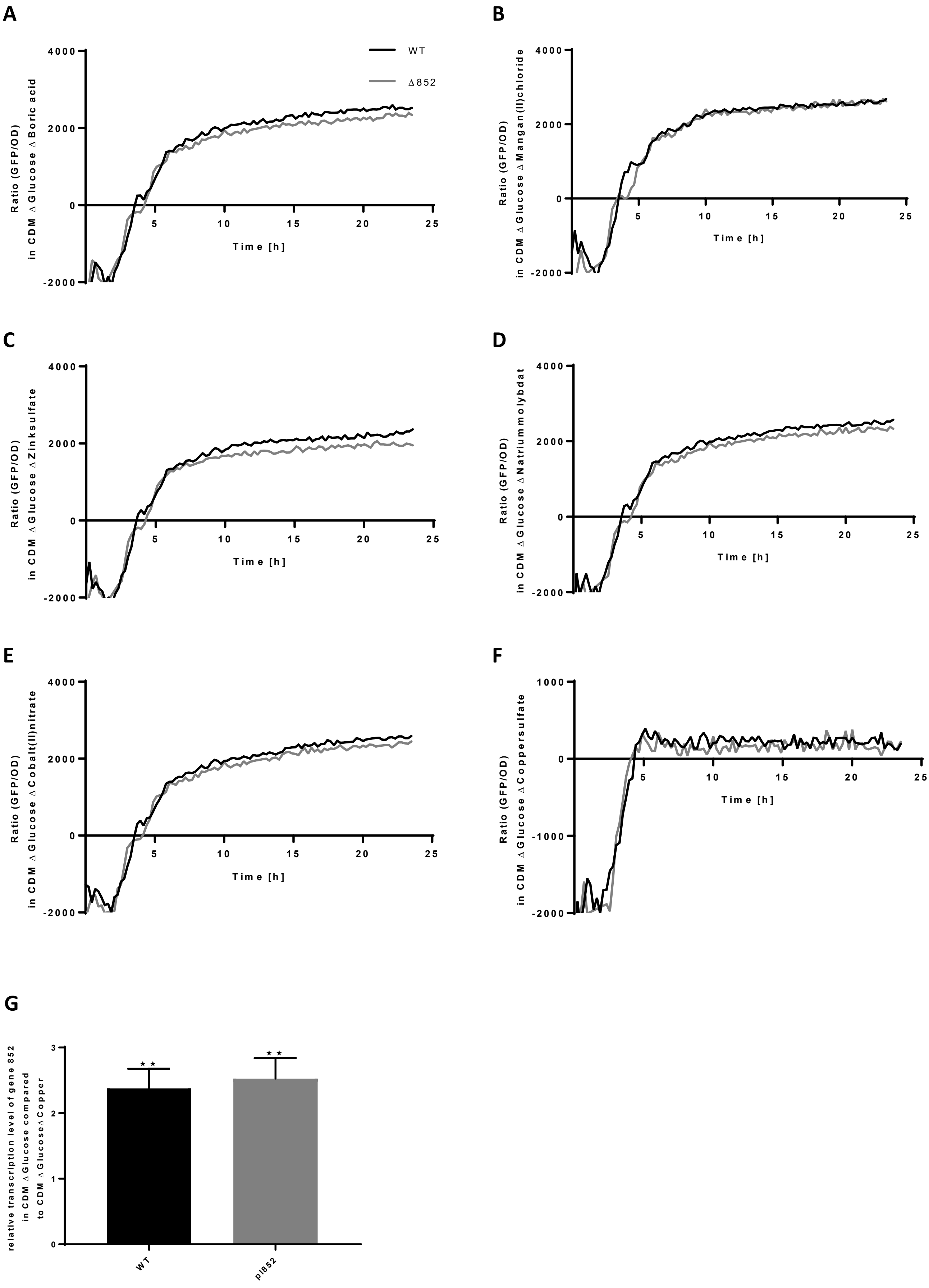

### Figure S3

#### Supplemental Figure 3

A

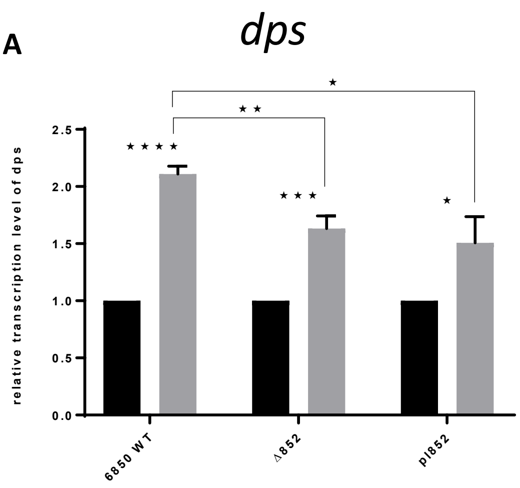


B

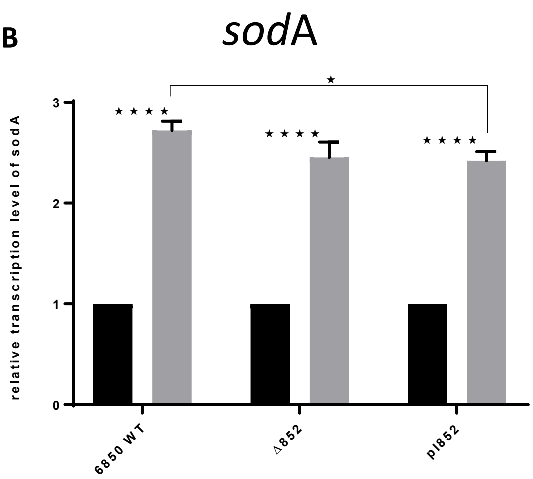

### Figure S4

#### Supplemental Figure 4

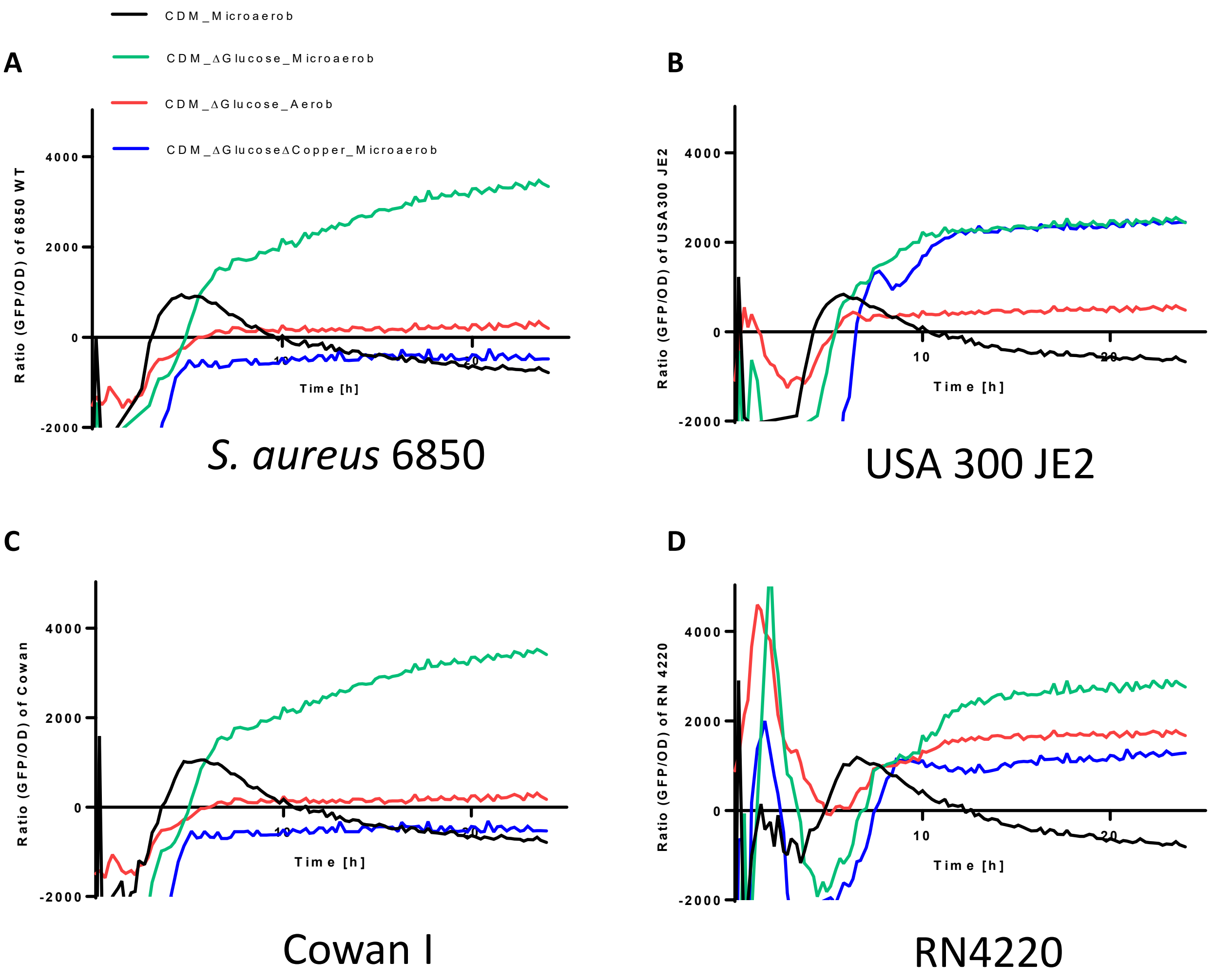
