## Supplementary material for "Identification of a novel LysR-type transcriptional regulator in *Staphylococcus aureus* that is crucial for secondary tissue colonization during metastatic bloodstream infection": Table S1

**Table S1: Bacterial strains and plasmids used in this study**

| Strain | Description | Source |
|---|---|---|
| *Escherichia coli* | | |
| DH5α | *fhuA2 lac(del)U169 phoA glnV44 Φ80' lacZ(del)M15 gyrA96 recA1 relA1 endA1 thi-1 hsdR17* | |
| *Staphylococcus aureus* | | |
| RN4220 | NCTC 8325-4 *sau1-, hsdR-,* laboratory strain accepting foreign DNA; β-toxin producer, non-hemolytic | (67) |
| 6850 | methicillin-sensitive; *spa* type t185, sequence type 50, Isolated from a patient with a skin abscess, progressed to bacteremia, osteomyelitis, septic arthritis, and multiple systemic abscesses | (30) |
| 6850 Δ852 | *S. aureus* 6850 with deletion of RSAU_000852 | This study |
| 6850 pl852 | *S. aureus* Δ852 complemented with pL852, Chloramphenicol-resistant | This study |

| Plasmid name | Description | Source |
|---|---|---|
| pl852 | Expression plasmid for RSAU_000852 under control of its native promoter | This study |
| p2085 | derivative of pALC2084 (68) | (20) |
| p2085-852 | plasmid for anhydrous tetracycline-inducible expression of RSAU_000852 | This study |
| p2085-Pr852-GFP | RSAU_000852 promoter fusion with green fluorescent protein (GFP) | This study |
