## Supplementary material for "Identification of a novel LysR-type transcriptional regulator in *Staphylococcus aureus* that is crucial for secondary tissue colonization during metastatic bloodstream infection": Table S2

### Table S2: Oligonucleotides used in this study

| Name | Sequence (5´-3´) |
|---|---|
| *RT-PCR* | |
| **RT-852-F** | TGAACAATATGAAAGTGGCA |
| **RT-852-R** | GGATGCTCGTTAAAGAATG |
| **RT-copA-F** | CGCTGTAGAAACTGTCGAATTAG |
| **RT-copA-R** | GGATAATAGTCAACTTTAGCTTGC |
| **RT-copB-F** | GTTGAAGGTATGAGCTGTGG |
| **RT-copB-R** | CGACATCGTAACCTTGATCTTC |
| **RT-gabT-F** | GTATCATGAACCGTTGGTACG |
| **RT-gabT-R** | GAGCCAAAAGTTGAACCATG |
| **RT-ilvA-F** | CATGTGCGAGTGCAGG |
| **RT-ilvA-R** | CAGTGATCAAATGTATCACCAGTG |
| **RT-ilvB-F** | GAACAAGGTGCTGTTCATGC |
| **RT-ilvB-R** | CTTGTCCGGTGAATACAACTAG |
| **RT-ilvC-F** | TGAAACATTAGTAGAAGCGG |
| **RT-ilvC-R** | CGTGGTCCTGAAACATAGTC |
| **RT-ilvD-F** | GGAATAGCTATGGGACATATCG |
| **RT-ilvD-R** | CAGAGCAAAAGATAGCTGGTAC |
| **RT-ilvH-F** | CCCGGGATTTCTAACATGG |
| **RT-ilvH-R** | GGCAGATGTTAACTAGCAGC |
| **RT-lacC-F** | GATCATGCCGGCATCAAG |
| **RT-lacC-R** | CCTGAAATAGCAACTGCTTC |
| **RT-lacE-F** | CTATACGATGGGGCTTGTAGC |
| **RT-lacE-R** | CCCATAAATGCACTTAAGAATCC |
| **RT-leuA2-F** | GGGATGGTATGCAACAAAGTAATGT |
| **RT-leuA2-R** | GGACTTAAAATGTCTCTGAATTGATG |
| **RT-leuB-F** | GCAATCGGTGGACCTAAATG |
| **RT-leuB-R** | CACGGACTATAACTAAATCTGTACC |
| **RT-leuC-F** | CGCCATAGATTTTGGGGTG |
| **RT-leuC-R** | GTTGCGAAAACATGTTCAACTTC |
| **RT-lrgA-F** | CTGGTGCTGTTAAGTTAGGC |
| **RT-lrgA-R** | GTATTGTTGAGACGATTATTAGTCC |
| **RT-msrA-F** | GGACCACGGTCTTGATATTG |
| **RT-msrA-R** | GGCGGACATATTGAAAATCC |
| **RT-purQ-F** | CTGAAGGTAAGCCAGTATTAGG |
| **RT-purQ-R** | CCGTGAGCTACAGGATATATAAC |
| **RT-pyrAB-F** | GAGCAACCTGACGCTTTAC |
| **RT-pyrAB-R** | GTTCTAAACATTTCACGGTCTTC |
| **RT-pyrB-F** | CGAATATTAACATCCCAATTGCG |
| **RT-pyrB-R** | GCACCTAATGCTTTTAAACTATGG |
| **RT-pyrC-F** | GACAATTGAAACTGGTACTAAAGC |
| **RT-pyrC-R** | CCTAATTGACGTGTTGTAATTGAAG |
| **RT-RNAIII-F** | ACATAGCACTGAGTCCAAGG |
| **RT-RNAIII-R** | TCGACACAGTGAACAAATTC |
| **RT-ureB-F** | CAGAGGTTGAAATTAATAACCAT |
| **RT-ureB-R** | CTCCAGCTGGAATATCTAAATG |
| **RT-ureC-F** | GCAGACCTTGTTATTTCTAATGC |
| **RT-ureC-R** | CAATACCACCAGCAGTGA |
| ***Cloning*** | |
| **852-BamHI-R** | TAATTGGATCCTTATAATTGTTCATTGGCAATATACTT |
| **852-NotI-F** | AAATAGCGGCCGCCGAACATTACTTTGTTGCATAC |
| **852_test_R** | TATCTCCATACAATTTCCAATC |
| **852_test_F** | AAGCGATTGATTTAGTTGAC |
| **attB2-852-down-R** | GGGGACCACTTTGTACAAGAAAGCTGGGTGTTTAAAT ATATTTTCACCAATTATAGGTTTG |
| **852-down-F-SacII** | GATCGACCGCGGGAACTTACCTCTTTCAAAAAAGTTAATAATT |
| **852-up-R-SacII** | GATCGACCGCGGAATAAATTTCAAATCTAAAAAACCAAGAATGC |
| **attB1-852-up-F** | GGGGACAAGTTTGTACAAAAAAGCAGGCTATAAACGTGTT GTAGGTCAAGATAAA |
| **pGFP-Inf-Prom F** | GAATTCTTAGGAGGATGATTATTTATGAGTAAAGGAGAAGAAC |
| **pGFP vec R** | GCATGCAAGCTTTTAAAAAGCAAATATGAGCCAAATAAA |
| **pGFP_852-Prom_fw** | TAAAAGCTTGCATGCGATATTTTGAAATAATTTC |
