## Supplementary material for "Identification of a novel LysR-type transcriptional regulator in *Staphylococcus aureus* that is crucial for secondary tissue colonization during metastatic bloodstream infection": Table S3

### Table S3: Enrichment in GO-terms as determined by STRING-DB.org

| GO-TERM (BIOLOGICAL PROCESS) | DESCRIPTION | FDR |
|---|---|---|
| **GO:0009082** | **branched-chain amino acid biosynthetic process** | **9.14e-06** |
| **GO:0009081** | branched-chain amino acid metabolic process | 9.14e-06 |
| **GO:1901607** | alpha-amino acid biosynthetic process | 9.45e-05 |
| **GO:1901605** | alpha-amino acid metabolic process | 9.45e-05 |
| **GO:0044281** | small molecule metabolic process | 9.45e-05 |
| **GO:0019627** | **urea metabolic process** | **9.45e-05** |
| **GO:0006520** | cellular amino acid metabolic process | 9.45e-05 |
| **GO:0008652** | cellular amino acid biosynthetic process | 0.00010 |
| **GO:0009097** | isoleucine biosynthetic process | 0.00031 |
| **GO:0006549** | isoleucine metabolic process | 0.00031 |
| **GO:1901564** | organonitrogen compound metabolic process | 0.0013 |
| **GO:0044283** | small molecule biosynthetic process | 0.0013 |
| **GO:0009987** | cellular process | 0.0020 |
| **GO:0008152** | metabolic process | 0.0024 |
| **GO:0006807** | nitrogen compound metabolic process | 0.0042 |
| **GO:0009099** | valine biosynthetic process | 0.0046 |
| **GO:0044205** | **'de novo' UMP biosynthetic process** | **0.0072** |
| **GO:0043419** | urea catabolic process | 0.0072 |
| **GO:0006573** | valine metabolic process | 0.0072 |
| **GO:1901566** | organonitrogen compound biosynthetic process | 0.0081 |
| **GO:0044237** | cellular metabolic process | 0.0081 |
| **GO:0009098** | leucine biosynthetic process | 0.0086 |
| **GO:0006551** | leucine metabolic process | 0.0086 |
| **GO:0071704** | organic substance metabolic process | 0.0111 |
| **GO:0046049** | UMP metabolic process | 0.0111 |
| **GO:0046132** | pyrimidine ribonucleoside biosynthetic process | 0.0191 |
| **GO:0046131** | pyrimidine ribonucleoside metabolic process | 0.0191 |
| **GO:0009156** | ribonucleoside monophosphate biosynthetic process | 0.0191 |
| **GO:0006812** | cation transport | 0.0253 |
| **GO:0009260** | ribonucleotide biosynthetic process | 0.0284 |
| **GO:0009161** | ribonucleoside monophosphate metabolic process | 0.0284 |
| **GO:0044238** | primary metabolic process | 0.0298 |
| **GO:0019637** | organophosphate metabolic process | 0.0327 |
| **GO:0019835** | cytolysis | 0.0339 |
| **GO:0044249** | cellular biosynthetic process | 0.0379 |
| **GO:0009259** | ribonucleotide metabolic process | 0.0392 |
| **GO:0009064** | glutamine family amino acid metabolic process | 0.0396 |
| **GO:0044248** | cellular catabolic process | 0.0412 |
