## Supplementary material for "Identification of a novel LysR-type transcriptional regulator in *Staphylococcus aureus* that is crucial for secondary tissue colonization during metastatic bloodstream infection": Table S4

**Table S4. Gene mutants identified enriched or depleted in murine liver after intravenous infection (cutoff: adj. p-Values <= 0.0005)**

| ID | $log_2FC$ | Adj. **p-Value** | gene | annotation |
|---|---|---|---|---|
| **RSAU_001097** | -5.74 | 0.0000233 | def | Peptide deformylase |
| **RSAU_000902** | -5.35 | 0.0000354 | ugtP | Processive diacylglycerol glucosyltransferase |
| **RSAU_000586** | -5.26 | 7.42E-07 | rsaC | non-coding RNA RsaC |
| **RSAU_000571** | -4.64 | 8.67E-08 | | Putative membrane protein |
| **RSAU_001052** | -3.57 | 0.0000599 | eta | C4-dicarboxylate transporter/malic acid transport family protein |
| **RSAU_000958** | -3.22 | 0.00002 | purM | Phosphoribosylformylglycinamidine cyclo-ligase |
| **RSAU_002404** | -3.19 | 0.0000223 | crtQ | 4,4'-diaponeurosporenoate glycosyltransferase |
| **RSAU_000985** | -3.02 | 0.000135 | potB | Spermidine/putrescine ABC transporter permease protein |
| **RSAU_002542** | -2.79 | 0.0000219 | | DNA-binding protein, putative |
| **RSAU_000940** | 3.44 | 0.000137 | atl | Bifunctional autolysin / N-acetylmuramoyl-L-alanine amidase |
| **RSAU_002096** | 3.74 | 5.78E-07 | pycA | RND multidrug transporter; Acriflavin resistance protein |
| **RSAU_001615** | 3.83 | 0.000134 | leuS | Leucyl-tRNA synthetase |
| **RSAU_001480** | 3.99 | 0.000131 | alaS | Alanyl-tRNA synthetase |
| **RSAU_000981** | 4.05 | 0.0000396 | pdhD | Dihydrolipoamide dehydrogenase |
| **RSAU_000014** | 4.06 | 0.0000659 | | Phosphoesterase, DHH family protein |
| **RSAU_002342** | 4.07 | 0.00002 | gntP | Gluconate permease |
| **RSAU_002402** | 4.18 | 0.000123 | crtN | Squalene/phytoene synthase |
| **RSAU_002410** | 4.21 | 0.000084 | isaA | Immunodominant antigen A, transglycosylase isaA |
| **RSAU_000222** | 4.22 | 0.00000207 | lytM | Peptidoglycan hydrolase |
| **RSAU_000494** | 4.22 | 0.00000151 | rpoB | DNA-directed RNA polymerase subunit beta |
| **RSAU_000758** | 4.25 | 0.0000938 | rnr | Ribonuclease R |
| **RSAU_001970** | 4.27 | 0.0000804 | | ATP-GRASP domain protein |
| **RSAU_002274** | 4.34 | 0.0000386 | bcr | Drug resistance transporter protein |
| **RSAU_001152** | 4.34 | 0.000135 | infB | Translation initiation factor IF-2 |
| **RSAU_002241** | 4.41 | 0.0000219 | fnt | Formate/nitrite transporter family protein |
| **RSAU_000888** | 4.41 | 0.000196 | | Monovalent cation/H+ antiporter-2 family protein |
| **RSAU_002255** | 4.5 | 0.000154 | gpmA | 2,3-bisphosphoglycerate-dependent phosphoglycerate mutase |
| **RSAU_001560** | 4.51 | 0.0000317 | dnaE | DNA polymerase III subunit alpha |
| **RSAU_000065** | 4.53 | 0.0000518 | | hypothetical protein |
| **RSAU_002302** | 4.62 | 0.0000219 | opp-1D | Oligopeptide ABC transporter, ATPase domain, putative |
| **RSAU_000266** | 4.63 | 0.000151 | geh | glycerol ester hydrolase |
| **RSAU_001210** | 4.63 | 0.0000451 | dhoM | Homoserine dehydrogenase |
| **RSAU_001882** | 4.68 | 0.0000219 | gcp | Putative DNA-binding/iron metalloprotein/AP endonuclease |
| **RSAU_000168** | 4.7 | 0.00018 | walK | Sensor histidine kinase protein |
| **RSAU_000943** | 4.75 | 0.0000599 | nanE | Cell envelope-related transcriptional attenuator domain protein |
| **RSAU_002525** | 4.83 | 0.00014 | hisZ | ATP phosphoribosyltransferase regulatory subunit |
| **RSAU_002298** | 4.89 | 0.000149 | gltB2 | Ferredoxin-dependent glutamate synthase |
| **RSAU_000666** | 4.91 | 0.0000599 | | Transporter Anion:Sodium Symporter family protein |
| **RSAU_001351** | 4.94 | 0.0000354 | ebpS | Elastin binding protein |
| **RSAU_002089** | 4.96 | 0.0000023 | pbuG | Xanthine/uracil permease family protein |
| **RSAU_002272** | 4.96 | 0.000123 | glxK | Glycerate kinase |

| | | | | |
|---|---|---|---|---|
| **RSAU_001253** | 4.98 | 0.00000748 | trpB | Tryptophan synthase beta chain |
| **RSAU_002290** | 4.99 | 0.000137 | | Amino acid transporter permease |
| **RSAU_000117** | 5.06 | 0.000125 | capO | Capsular polysaccharide synthesis UDP-N-acetyl-D dehydrogenase |
| **RSAU_000844** | 5.13 | 0.00000323 | addA | ATP-dependent helicase/nuclease subunit A |
| **RSAU_002453** | 5.13 | 0.0000023 | gbsA | Glycine betaine aldehyde dehydrogenase |
| **RSAU_001518** | 5.14 | 0.000166 | | hypothetical protein |
| **RSAU_000906** | 5.15 | 0.000193 | terC | Toxic anion resistance membrane protein |
| **RSAU_000937** | 5.15 | 0.0000712 | aspC | Putative aminotransferase |
| **RSAU_001219** | 5.18 | 0.000052 | guaC | Guanosine monophosphate reductase |
| **RSAU_000592** | 5.19 | 0.000123 | tagA | Teichoic acid biosynthesis protein |
| **RSAU_000600** | 5.19 | 0.000162 | nupC2 | Na+ dependent nucleoside transporter protein |
| **RSAU_001943** | 5.19 | 0.000135 | atpA | FOF1 ATP synthase subunit alpha |
| **RSAU_000994** | 5.22 | 5.78E-07 | typA | GTP-binding protein TypA/BipA |
| **RSAU_002485** | 5.22 | 0.0000599 | pmi | Mannose-6-phosphate isomerase |
| **RSAU_001860** | 5.24 | 0.00000798 | groEL | 60 kDa chaperonin GroEL |
| **RSAU_002334** | 5.24 | 0.0000451 | pgcA | Phosphoglucomutase/phosphomannomutase |
| **RSAU_000265** | 5.28 | 0.0000283 | sirA | Nucleoside recognition domain protein |
| **RSAU_002372** | 5.32 | 0.000156 | sdaAA | L-serine dehydratase, iron-sulfur-dependent, alpha subunit |
| **RSAU_002291** | 5.34 | 0.0000668 | pnbA | Para-nitrobenzyl esterase chain A |
| **RSAU_000450** | 5.36 | 0.000123 | ftsH | Cell division protein FtsH, putative |
| **RSAU_000794** | 5.37 | 0.0000651 | sufS | Cysteine desulfurase, SufS subfamily |
| **RSAU_001157** | 5.39 | 0.00000436 | pnpA | Polyribonucleotide nucleotidyltransferase |
| **RSAU_002278** | 5.45 | 0.000162 | cycA | Amino acid permease, putative |
| **RSAU_002357** | 5.45 | 0.00013 | fbp | Fructose-1,6-bisphosphatase class 3 |
| **RSAU_001388** | 5.49 | 0.0000124 | ispA | Geranyltranstransferase |
| **RSAU_000974** | 5.51 | 0.000032 | rnjA | Ribonuclease J 1 |
| **RSAU_001991** | 5.55 | 0.0000451 | glmS | Glucosamine--fructose-6-phosphate aminotransferase |
| **RSAU_002484** | 5.65 | 0.0000389 | manP | PTS system, fructose-specific IIABC component |
| **RSAU_002100** | 5.65 | 0.000128 | | Major facilitator transporter |
| **RSAU_000062** | 5.69 | 0.00000243 | | 67 kDa myosin-cross-reactive antigen |
| **RSAU_001080** | 5.69 | 0.0000932 | rluD | Ribosomal large subunit pseudouridine synthase |
| **RSAU_001786** | 5.71 | 0.00000788 | gatB | Aspartyl/glutamyl-tRNA amidotransferase subunit B |
| **RSAU_000693** | 5.72 | 0.0000599 | ltaS | Glycerol phosphate lipoteichoic acid synthase |
| **RSAU_002444** | 5.72 | 0.00000207 | | Amino acid permease family protein |
| **RSAU_000739** | 5.75 | 0.0000733 | trxB | Thioredoxin reductase |
| **RSAU_001145** | 5.82 | 0.0000497 | proS | Prolyl-tRNA synthetase |
| **RSAU_000718** | 5.86 | 7.42E-07 | pepT | Peptidase T |
| **RSAU_000864** | 5.88 | 0.000117 | oppA | Oligopeptide ABC transporter-binding protein OppA |
| **RSAU_000833** | 5.9 | 0.000135 | cls | Family NADH-dependent flavin oxidoreductase, putative |
| **RSAU_002020** | 5.96 | 0.0000606 | htsB | Iron ABC transporter, permease protein |
| **RSAU_000289** | 5.99 | 0.00000791 | efeB | Dyp-type peroxidase family protein |
| **RSAU_002289** | 6 | 0.00000748 | rpsP | Bacteriocin-protection, YdeI/OmpD-Associated family protein |
| **RSAU_002442** | 6.05 | 0.000137 | budA | Alpha-acetolactate decarboxylase |
| **RSAU_000420** | 6.12 | 0.000159 | ribA | Protein from nitrogen regulatory protein P-II family YAAQ |
| **RSAU_001032** | 6.12 | 0.00000436 | sdhA | Succinate dehydrogenase flavoprotein subunit |
| **RSAU_001394** | 6.14 | 0.0000233 | accB | Biotin carboxyl carrier protein of acetyl-CoA carboxylase |

| | | | | |
|---|---|---|---|---|
| **RSAU_002041** | 6.14 | 0.0000466 | cobB | NAD-dependent protein deacetylase SIR2 family |
| **RSAU_001189** | 6.18 | 0.000117 | hflX | GTP-binding protein HflX |
| **RSAU_002276** | 6.31 | 5.91E-07 | | Peptidase C39 like family protein |
| **RSAU_002432** | 6.32 | 0.000126 | | Adenine nucleotide alpha hydrolases superfamily protein |
| **RSAU_000300** | 6.47 | 0.0000194 | | Cyclase family protein |
| **RSAU_002165** | 6.5 | 3.46E-07 | hipO2 | N-L-amino acid amidohydrolase |
| **RSAU_000093** | 6.54 | 0.0000219 | drm | Phosphopentomutase |
| **RSAU_000497** | 6.56 | 0.000195 | rpsL | 30S ribosomal protein S12 |
| **RSAU_001643** | 6.56 | 0.0000213 | metK | S-adenosylmethionine synthetase |
| **RSAU_001034** | 6.69 | 0.0000451 | murI | Glutamate racemase |
| **RSAU_000566** | 6.84 | 0.0000314 | btuF | Iron ABC transporter binding protein |
| **RSAU_000534** | 6.92 | 0.000117 | | putative Membrane protein |
| **RSAU_000505** | 7 | 0.0000147 | capD | NAD dependent epimerase/dehydratase family protein |
| **RSAU_000506** | 7.03 | 0.0000354 | ilvE | Branched-chain amino acid aminotransferase |
| **RSAU_001527** | 7.03 | 0.0000512 | hemB | Delta-aminolevulinic acid dehydratase |
| **RSAU_000498** | 7.25 | 0.000117 | rpsG | 30S ribosomal protein S7 |
| **RSAU_000427** | 7.38 | 0.00000243 | metG | Methionyl-tRNA synthetase |
| **RSAU_002292** | 7.91 | 1.21E-09 | rimM | Major facilitator transporter, chloramphenicol |
