## Supplementary material for "Identification of a novel LysR-type transcriptional regulator in *Staphylococcus aureus* that is crucial for secondary tissue colonization during metastatic bloodstream infection": Table S5

**Table S5: *S. aureus* transposon mutants found either enriched or depleted in both, kidney or liver of mice after intravenous infection by TN-seq.** Counts of sequenced transposon insertion sites were compared with the counts obtained from the infection inoculum. By comparing this inocolum to the recovered samples genes which are lost in a specific tissue are represented by negative $log2_FC$ and vice versa.

| 6850 Locus ID | NCTC8325 Locus ID | Gene | Annotation | log2FC Inoc./ Kidney | Adj. p-Value Kidney (<1e-3) | logFC Inoc./ Liver | Adj. p-Value Liver (<1e-3) |
|---|---|---|---|---|---|---|---|
| **RSAU_000958** | SAOUHSC_01015 | purM | Phosphoribosylformyl glycinamidine cyclo-ligase | -5 | 0.00015 | -3.22 | 0.00002 |
| **RSAU_000571** | SAOUHSC_00618 | | Putative membrane protein | -4.04 | 0.000124 | -4.64 | 8.67E-08 |
| **RSAU_002542** | SAOUHSC_03032 | | DNA-binding protein, putative | -2.94 | 0.000124 | -2.79 | 0.0000219 |
| **RSAU_000494** | SAOUHSC_00524 | rpoB | DNA-directed RNA polymerase subunit beta | 3.72 | 0.000275 | 4.22 | 0.00000151 |
| **RSAU_000940** | SAOUHSC_00994 | atl | Bifunctional autolysin / N-acetylmuramoyl-L-alanine amidase / Endo-N | 3.74 | 0.00024 | 3.44 | 0.000137 |
| **RSAU_000222** | SAOUHSC_00248 | lytM | Peptidoglycan hydrolase | 3.96 | 0.000124 | 4.22 | 0.00000207 |
| **RSAU_000981** | SAOUHSC_01043 | pdhD | Dihydrolipoamide dehydrogenase | 4.09 | 0.000285 | 4.05 | 0.0000396 |
| **RSAU_001882** | SAOUHSC_02277 | gcp | Putative DNA-binding/iron metalloprotein/AP endonuclease | 4.26 | 0.000388 | 4.68 | 0.0000219 |
| **RSAU_001970** | SAOUHSC_02373 | | ATP-grasp domain protein | 4.4 | 0.000262 | 4.27 | 0.0000804 |
| **RSAU_002089** | SAOUHSC_02516 | pbuG | Xanthine/uracil permease family protein | 4.43 | 0.000202 | 4.96 | 0.0000023 |
| **RSAU_000495** | SAOUHSC_00525 | rpoC | DNA-directed RNA polymerase subunit beta' | 4.5 | 0.000774 | 4.41 | 0.000206 |
| **RSAU_000943** | SAOUHSC_00997 | nanE | Cell envelope-related transcriptional attenuator domain protein | 4.53 | 0.000632 | 4.75 | 0.0000599 |
| **RSAU_002274** | SAOUHSC_02725 | bcr | Drug resistance transporter protein | 4.53 | 0.000189 | 4.34 | 0.0000386 |
| **RSAU_000822** | SAOUHSC_00882 | | Pyridine nucleotide-disulfide oxidoreductase family protein | 4.57 | 0.000291 | 3.95 | 0.000453 |
| **RSAU_001560** | SAOUHSC_01811 | dnaE | DNA polymerase III subunit alpha | 4.81 | 0.000124 | 4.51 | 0.0000317 |
| **RSAU_000844** | SAOUHSC_00905 | addA | ATP-dependent helicase/nuclease subunit A | 4.84 | 0.000124 | 5.13 | 0.00000323 |
| **RSAU_002334** | SAOUHSC_02793 | pgcA | Phosphoglucomutase / phosphomannomutase | 5 | 0.000783 | 5.24 | 0.0000451 |
| **RSAU_000735** | SAOUHSC_00781 | hprK | HPr kinase/phosphorylase | 5.04 | 0.00081 | 4.64 | 0.000453 |
| **RSAU_000193** | SAOUHSC_00217 | gutB | Zinc-binding sorbitol dehydrogenase | 5.1 | 0.000236 | 4.2 | 0.000667 |
| **RSAU_001351** | SAOUHSC_01501 | ebpS | Elastin binding protein | 5.14 | 0.000124 | 4.94 | 0.0000354 |
| **RSAU_002484** | SAOUHSC_02975 | manP | PTS system, fructose-specific IIABC component | 5.15 | 0.000555 | 5.65 | 0.0000389 |
| **RSAU_002525** | SAOUHSC_03015 | hisZ | ATP phosphoribosyltransferase regulatory subunit | 5.16 | 0.000305 | 4.83 | 0.00014 |
| **RSAU_000259** | SAOUHSC_00293 | nupC | Nucleoside transporter permease | 5.17 | 0.00092 | 5 | 0.000229 |
| **RSAU_000009** | SAOUHSC_00009 | serS | Seryl-tRNA synthetase | 5.19 | 0.000124 | 4.07 | 0.000819 |
| **RSAU_000666** | SAOUHSC_00698 | | Transporter Anion:Sodium Symporter family protein | 5.22 | 0.00015 | 4.91 | 0.0000599 |
| **RSAU_001219** | SAOUHSC_01330 | guaC | Guanosine monophosphate reductase | 5.28 | 0.00025 | 5.18 | 0.000052 |
| **RSAU_000600** | SAOUHSC_00648 | nupC2 | Na+ dependent nucleoside transporter protein | 5.31 | 0.000774 | 5.19 | 0.000162 |
| **RSAU_002444** | SAOUHSC_02923 | | Amino acid permease family protein | 5.35 | 0.000117 | 5.72 | 0.00000207 |
| **RSAU_000994** | SAOUHSC_01058 | typA | GTP-binding protein TypA/BipA | 5.58 | 0.000000408 | 5.22 | 0.000000578 |

| | | | | | | | |
|---|---|---|---|---|---|---|---|
| **RSAU_002289** | SAOUHSC_02747 | rpsP | Bacteriocin-protection, YdeI/OmpD-Associated family protein | 5.6 | 0.000196 | 6 | 0.00000748 |
| **RSAU_001080** | SAOUHSC_01163 | rluD | Ribosomal large subunit pseudouridine synthase | 5.62 | 0.000742 | 5.69 | 0.0000932 |
| **RSAU_002278** | SAOUHSC_02729 | cycA | Amino acid permease, putative | 5.7 | 0.000486 | 5.45 | 0.000162 |
| **RSAU_000062** | SAOUHSC_00061 | | 67 kDa myosin-cross-reactive antigen | 5.76 | 0.0000324 | 5.69 | 0.00000243 |
| **RSAU_001145** | SAOUHSC_01240 | proS | Prolyl-tRNA synthetase | 5.77 | 0.000291 | 5.82 | 0.0000497 |
| **RSAU_000638** | SAOUHSC_00668 | vraG | ABC transporter permease | 5.94 | 0.000124 | 5.02 | 0.000259 |
| **RSAU_000864** | SAOUHSC_00927 | oppA | Oligopeptide ABC transporter-binding protein OppA | 5.97 | 0.000422 | 5.88 | 0.000117 |
| **RSAU_002020** | SAOUHSC_02428 | htsB | Iron ABC transporter, permease protein | 6.21 | 0.000202 | 5.96 | 0.0000606 |
| **RSAU_000696** | SAOUHSC_00731 | | Glycine betaine/choline ABC transporter ATP-binding protein | 6.23 | 0.000521 | 5.48 | 0.000688 |
| **RSAU_000427** | SAOUHSC_00461 | metG | Methionyl-tRNA synthetase | 6.5 | 0.00024 | 7.38 | 0.00000243 |
| **RSAU_002432** | SAOUHSC_02911 | | Adenine nucleotide alpha hydrolases superfamily protein | 6.6 | 0.000309 | 6.32 | 0.000126 |
| **RSAU_002292** | SAOUHSC_02752 | rimM | Major facilitator transporter, chloramphenicol | 7.11 | 0.000000322 | 7.91 | 1.21E-09 |
| **RSAU_000505** | SAOUHSC_00535 | capD | NAD dependent epimerase/dehydratase family protein | 7.2 | 0.0000655 | 7 | 0.0000147 |
